# Decoding the Transcriptome Dark Matter: Construction of Single-Cell Whole-Transcriptome Regulatory Atlas by dropTotal

**DOI:** 10.64898/2026.08.25.747146

**Authors:** Xudong Liu, Wenjian Cao, Yating Pan, Ziqing Luo, Tianhao Wu, Yupeng Du, Xidong Xu, Zhiguo Jin, Chengqi Li, Ying Mu, Yingchao Liu, Qiangyuan Zhu

**Author notes:** Corresponding author. (YM); (YL); (QZ). These authors contributed equally: Xudong Liu, Wenjian Cao and Yating Pan.

## Abstract

To profile unknown ncRNAs—“dark matter” in single cells, we developed dropTotal, a high-throughput droplet-based total RNA-seq method that uses dU-modified GAT primer with temperature-ramp hybridization and droplet merge barcoding to co-detect coding and non-coding transcripts with record sensitivity (>13,500 genes/cell, including >2,000 lncRNAs and >500 sncRNAs), compatible with fresh, frozen, fixed, and FFPE tissues. Applied to ∼75,000 human glioma nuclei, it captured 60,313 genes (18,681 lncRNA, 19,859 mRNAs and 6,753 sncRNAs), enabling ncRNA-driven regulatory landscape construction. In oligodendroglioma, module analysis identified recurrence-associated ncRNA-centered modules linked to therapy resistance and invasion; in glioblastoma, six cellular states showed hundreds of state-specific unannotated ncRNAs with divergent functions, from *MIR222HG*-mediated immune modulation to *SCIRT*-driven hypoxia adaptation. Alternative splicing analysis identified 428 state-specific junction markers and mapped cell-state-specific alternative splicing regulation. dropTotal offers broad application for decoding the underlying ncRNA biology and single-cell whole transcriptome regulatory mechanisms in cellular identity and disease progression.

## Introduction

The eukaryotic transcriptome consists of a broad spectrum of transcripts, including messenger RNAs (mRNAs) that code for proteins, and non-coding RNAs (ncRNAs) that function in gene regulation and other cellular processes^1,2^. In contrast to mRNAs, a great number of unknown ncRNAs—the predominant transcriptional output of the genome—constitute essential “dark matter” within these networks, forming intricate interconnections with other regulators to sculpt the transcriptome. Understanding these mechanisms requires knowledge of when and where genes are co-expressed. Single-cell RNA sequencing (scRNA-seq) enables this but is fundamentally limited by its reliance on poly(A) tail capture (Fig. 1a), which fails to detect the vast landscape of non-polyadenylated ncRNAs that contribute to regulatory mechanisms underlying differential mRNA expression^3–8^. This gap prevents the construction of complete co-expression relationships and the systematic dissection of critical ncRNA-target RNA interactions. Thus, despite advances in single-cell transcriptomics, deciphering this regulatory “dark matter”—the non-coding transcriptome—remains at an early stage.

**Fig. 1.**
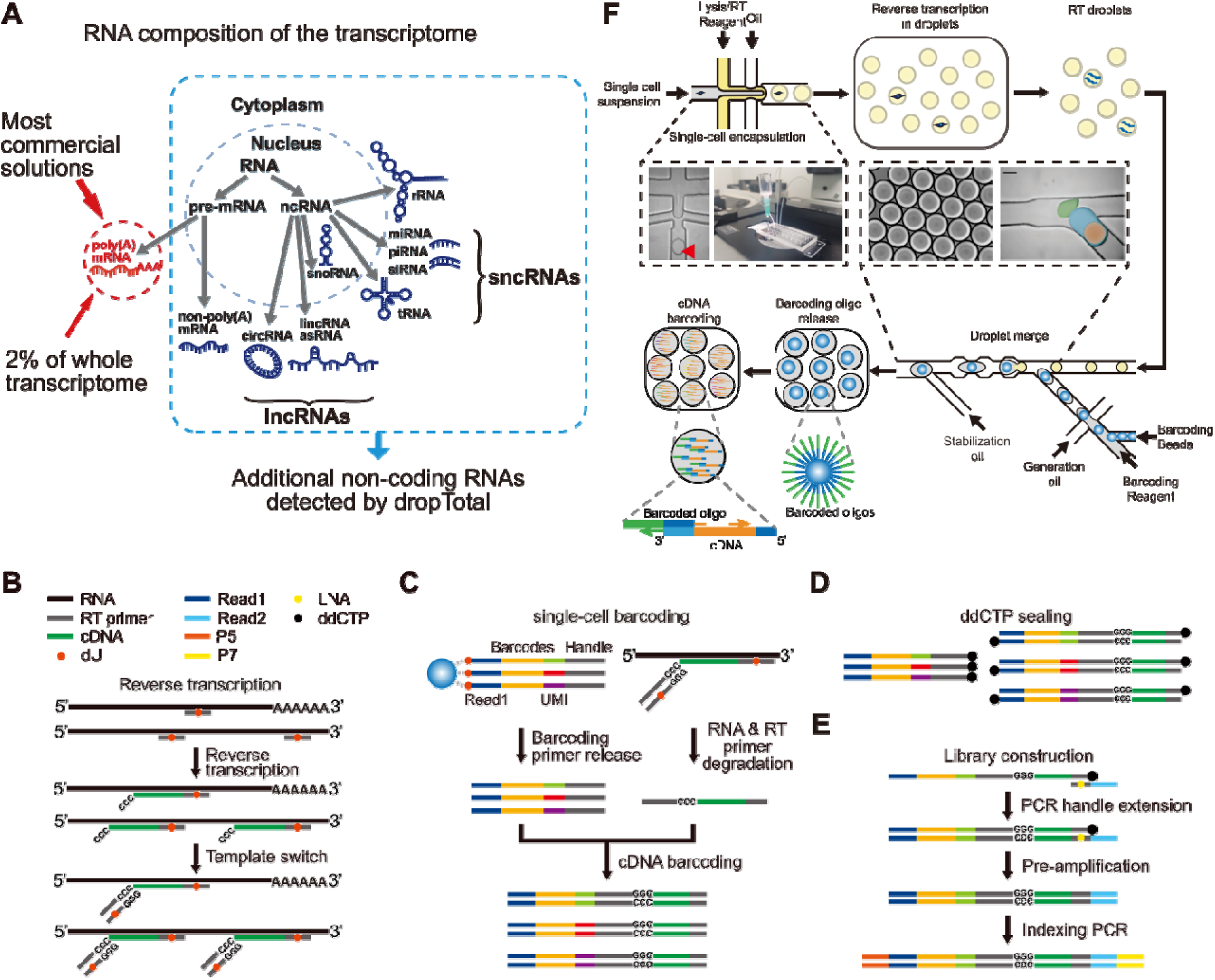
Microfluidic and chemical workflow of dropTotal. (A) Challenges in whole transcriptome detection by major high-throughput single-cell transcriptome methods. (B-E) Chemistry overview of dropTotal. (F) Microfluidic workflow of dropTotal. RT/lysis mix and single cells are co-encapsulated into droplets. After RT reaction, the RT droplets are merged with droplets containing barcoding reaction mix and BHBs for single-cell barcoding.

In this work, we developed dropTotal, a novel high-throughput droplet-based single-cell total RNA-seq method for coding and non-coding RNA profiling and comprehensive regulatory landscape construction. The core feature of dropTotal lies in the use of an optimally designed universal primer composed of the three bases G, A and T to efficiently capture coding and non-coding RNAs in single cells under a gradually increasing temperature, the generation of asymmetric cDNA products via deoxyuridine (dU) base modification and digestion of the RT primer for highly efficient single-cell barcoding, and a robust droplet-merging microfluidic platform that achieves high throughput single-cell and barcoding beads co-encapsulation by controlling changes in surfactant concentration and surface tension. Benchmarking against all of the high-throughput methods demonstrates its sensitive detection of a broad spectrum of coding and non-coding RNAs. By applying dropTotal to ∼75,000 nuclei from human frozen glioma samples, we generated cell-type-resolved expression atlases of ncRNAs, including long non-coding RNAs (lncRNAs), miscellaneous RNAs (miscRNAs), small nuclear RNAs (snRNAs), microRNAs (miRNAs), small nucleolar RNAs (snoRNAs), and revealed previously obscured layers of transcriptomic heterogeneity^9–11^. This enhanced detection enabled the identification of ncRNA-driven gene modules that distinguish primary from recurrent oligodendroglioma (OG), pinpointing key drivers of tumor progression. Furthermore, dropTotal enabled systematic dissection of the divergent regulatory functions exerted by ncRNAs across six distinct cellular states in glioblastoma (GBM). We further delineated the resulting cellular-state-specific alternative splicing landscape mediated by these ncRNAs. Collectively, our findings establish dropTotal as a powerful and broadly applicable tool for probing the full spectrum of transcriptional regulation in single cells. While exemplified here in glioma, the high-resolution atlases and regulatory principles defined by this approach provide a foundational resource and framework for understanding how ncRNAs orchestrate cellular identity and dynamics across diverse biological contexts, including development, homeostasis, and disease.

## Results

### dropTotal enables high-throughput detection of both non-coding and coding transcripts in single cells

Strategies for whole-transcriptome capture in single-cell sequencing can be broadly divided into two categories. The first, represented by VASA-drop^12^, relies on high temperature RNA fragmentation followed by poly(A) tailing, so that fragmented RNAs can be captured using poly(T) primers. However, this fragmentation approach based on high temperature inevitably introduces contamination from residual gDNA and RNA loss, generating substantial strand-specific noise^13^. The second, represented by MATQ-drop^14^, captures the entire transcriptome by employing MALBAC primers that anneal to arbitrary positions along the transcripts. To achieve efficient annealing of the MALBAC primers, a low-temperature step is required to promote non-specific binding between the primers and RNA; this low-temperature process gives rise to extensive primer-primer crosstalk and decreases detection sensitivity. To address the above limitations and enable the detection of both non-coding and coding transcripts at single-cell resolution, we first introduced an optimized universal primer. The universal primer contains only G, A and T bases (GAT primer) to achieve highly efficient and uniform annealing with RNA sequences and full-gene-body coverage during reverse transcription (RT)^15^, eliminating the issue of primer crosstalk. Three consecutive G bases at the 3’ end allow the universal primer to serve as template switching oligo (TSO) for barcoding handle addition (Fig. 1b).

The 5’ and 3’ ends of the cDNA generated by RT are reverse-complementary due to single primer design, which compromises the efficiency of the single-cell barcoding reaction. To allow unidirectional barcoding of cDNAs, elimination of potential crosstalk in downstream amplification as well as removal of excessive GAT primers and primer dimers, we introduced a deoxyuridine (dU) base in the middle of GAT primer. The 5’ ends of the barcoded oligonucleotides on the barcoded hydrogel beads (BHBs) were also modified with dU. With the presence of USER enzyme in barcoding reagent, dU bases are removed to release barcoded oligos, degrade excessive GAT primers and break the symmetry of cDNAs (Fig. 1c). Meanwhile, RNAs from single cells are degraded by RNaseH and RNaseIf, releasing cDNAs as templates for barcoded second-strand synthesis. Barcoded oligos hybridize to the full-length reverse-complementary GAT primer at the 3’ end of cDNAs to complete second-strand synthesis. Subsequently, the barcoded second-strand products are subjected to size selection to remove the majority of unused barcoded oligonucleotides. Any remaining unused barcoded oligonucleotides are sealed by ddCTP to prevent potential cell barcode swapping in the downstream library preparation (Fig. 1d).

To amplify barcoded second-strand products, an extension reaction is performed to add Illumina Read2 sequence. Since the remaining GAT primer sequence at the 3’ end of barcoded second-strand products is only 14 nt in length, three locked nucleic acid (LNA) bases are introduced in the Read2 extension primer and multiple cycles of annealing are performed to achieve high extension efficiency. The converted products further undergo pre-amplification and indexing PCR to generate complete libraries for sequencing (Fig. 1e).

To extend the single-cell whole-transcriptome sequencing technology described above to high throughput, we developed a robust active droplet merging platform (Fig. 1f and Extended Data Fig. 1). This novel droplet-merging platform employs a two-step reaction strategy to overcome the incompatibility between the RT and single-cell barcoding reactions. These reactions require distinct buffer and temperature conditions and therefore cannot be performed in the same reaction vessel. By controlling changes in surfactant concentration and interfacial tension alteration, the platform enables precise merging of droplets containing RT products with droplets encapsulating barcoding reagents and BHBs. Compared with electrode-based merge approaches, this tension-driven active merge eliminates the need for additional electrodes or microstructures on the chip, simplifying device design and achieving more stable droplet merge, thereby providing a solid technical foundation for high-throughput single-cell whole-transcriptome sequencing.

### Sensitive detection of both non-coding and coding transcripts by dropTotal

To benchmark dropTotal, we performed a species-mixing experiment with human HEK293T cells and mouse NIH3T3 cells. As shown in Fig. 2a, we identified 214 high quality cell barcodes. Based on species specificity, these cell barcodes were assigned to 132 human cells, 80 mouse cells and 2 collision events, showing a low doublet rate of 0.93% (Fig. 2b). High species specificity of unique molecular identifiers (UMIs) was observed for both species, 99.37% for human cells and 97.67% for mouse cells (Fig. 2c). At an average sequencing depth of 890,000 trimmed reads per cell, dropTotal captured a median of 68,661 UMIs and 13,521 genes for 293T cells, and a median of 40,659.5 UMIs and 8,242 genes for NIH3T3 cells (Fig. 2d,e).

**Fig. 2.**
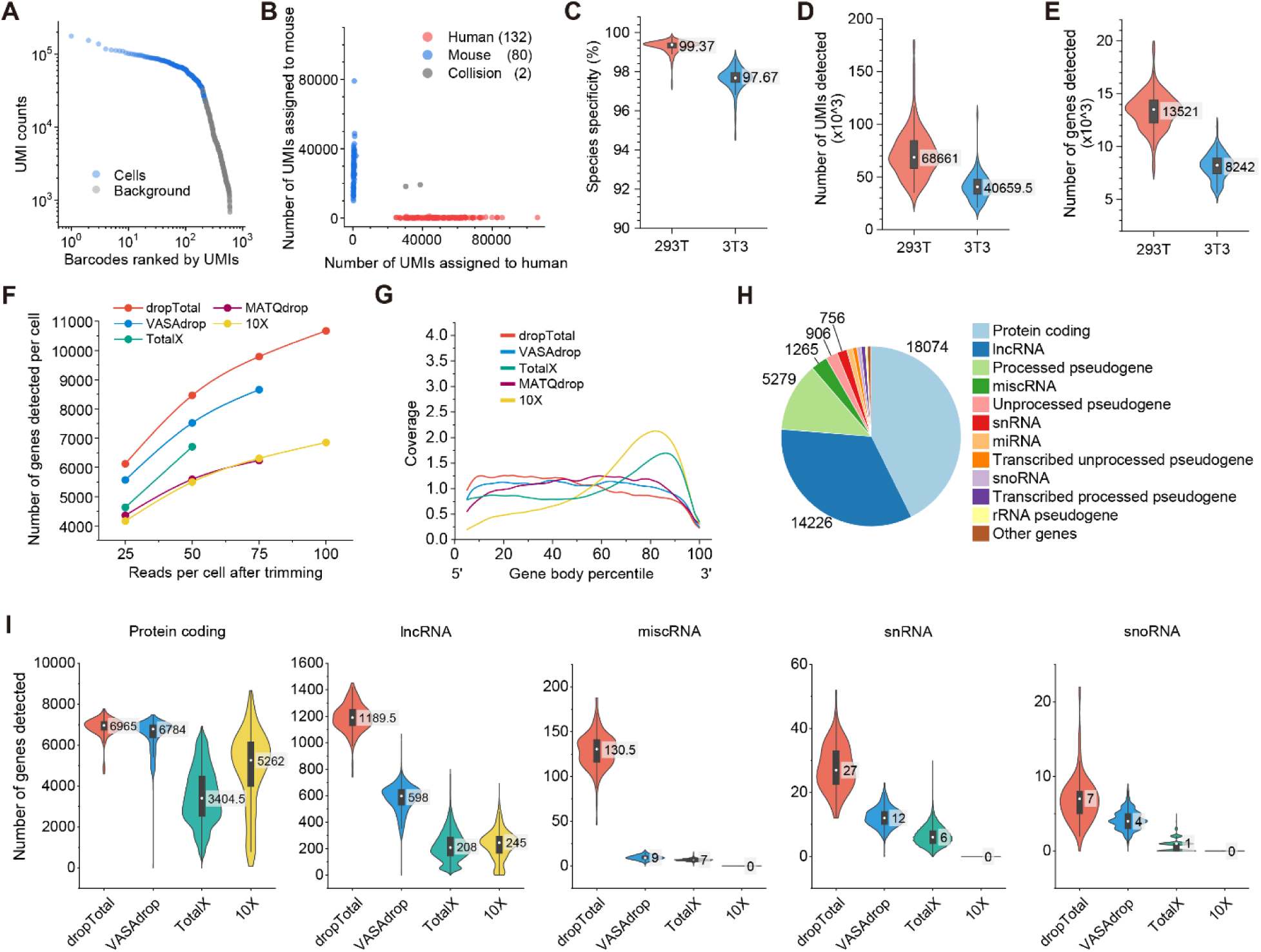
Sensitive detection of coding and non-coding transcripts by dropTotal. (A) UMI-barcode plot of cell line mixing experiment. Top 600 barcodes were shown. (B) Composition of annotated UMIs of 214 cell barcodes identified. (C) Species specificity of UMIs. (D and E) Detection sensitivity of dropTotal in UMI counts (D) and gene counts (E). (F) Number of genes detected plotted against reads used for analysis of high-throughput methods. (G) Gene body coverage of dropTotal and other methods. (H) Composition of gene types detected in HEK293T cells. (I) Number of protein coding, lncRNA and sncRNA (including miscRNA, snRNA and snoRNA) genes detected in HEK293T cells by dropTotal, VASA-drop, TotalX and 10X at depth of 50k reads.

We then compared dropTotal to the following methods: widely used commercial platform, 10x Chromium as well as 3 high-throughput single cell methods with total RNA detection capability, MATQ-drop^14^, VASA-seq^12^ and TotalX^16^. HEK293T datasets were downsampled for a fair comparison of gene detection sensitivity and saturation evaluation (Fig. 2f). dropTotal exhibited highest number of gene detection, with a median of 10,671 detected genes per cell, respectively, at a sequencing depth of 100,000 reads per cell (Fig. 2f). At the sequencing depth of 50,000 reads per cell, dropTotal showed higher gene detection rate (8,463 detected genes per cell) than VASA-drop (7,521 detected genes per cell). Meanwhile, dropTotal outperformed TotalX (6,706 detected genes per cell), MATQ-drop (5,593 detected genes per cell), 10x Chromium (5,498 detected genes per cell, Fig. 2f).

Owing to its efficient and uniform hybridization strategy, dropTotal provided even coverage across gene bodies. dropTotal, VASA-drop, and MATQ-drop showed relatively uniform gene-body coverage, with dropTotal exhibiting a slight 5’-end bias and MATQ-drop a slight 3’-end bias (Fig. 2g). TotalX uses a poly(A)-dependent barcoding strategy similar to that of 10x Chromium and therefore exhibited a pronounced 3′-end bias (Fig. 2g).

Poly(A)-tail independent chemistry enables dropTotal to detect a variety type of transcripts effectively. As shown in Fig. 2h, protein-coding genes consists 42.7% of all genes detected by dropTotal, followed by lncRNAs (33.6%) and processed pseudogenes (12.5%), which also participate in regulation of gene expression. The capability of dropTotal to capture a broad spectrum of non-coding RNAs is crucial to reveal their regulatory functions in biological processes. We compared the number of detected protein-coding, long non-coding and short non-coding genes by dropTotal, VASA-drop and 10x Chromium at the sequencing depth of 50,000 trimmed reads per cell (Fig. 2i). dropTotal detected more protein-coding genes (with a median of 6,965) than VASA-drop (with a median of 6,784), outperforming TotalX (with a median of 6,239) and 10x Chromium (with a median of 6,019). What’s more, for lncRNAs, dropTotal detected a median of 1,189.5 genes, nearly 2-fold of the genes detected by VASA-drop (with a median of 598). Although TotalX incorporated poly(A) tailing of RNAs, it only detected a median of 413 lncRNA genes. While 10x Chromium only detected a median of 245.5 lncRNAs. dropTotal also detected more sncRNAs than both VASA-drop and TotalX, while 10x Chromium is unable to detect any sncRNAs (Fig. 2i).

Overall, by combining a novel droplet merging microfluidic platform, shotgun and uniform hybridization by universal RT primer as well as dU-based primer degradation and symmetry breaking strategy, dropTotal achieved high-throughput and highly sensitive profiling of both non-coding and coding transcripts at single-cell resolution. Even coverage across the gene body allowed dropTotal to capture splicing junctions for further alternative splicing analyses.

### Cell-type-resolved profiling of non-coding RNAs in human frozen glioma samples

We next leveraged the transcriptome-wide capture efficiency of dropTotal and applied it to frozen clinical glioma samples to delineate the coding and non-coding landscapes of tumor and normal cells. Benefiting from the high capture efficiency of dropTotal across the transcriptome, we identified a total of 60,313 distinct genes in these glioma samples. Among them, 19,859 were mRNAs, accounting for approximately 32.9%. Notably, lncRNAs numbered 18,681, representing roughly 31.0% of the detected genes—a proportion nearly on par with that of mRNAs. In addition, we detected a broad spectrum of sncRNAs, including 2,196 miscRNAs, 1,867 snRNAs, 1,764 miRNAs, and 926 snoRNAs, collectively amounting to 6,753 species (approximately 11.2%) (Fig. 3a). These findings collectively demonstrate the robust capability of dropTotal to efficiently capture non-coding RNAs from frozen clinical samples. At the single-cell level, with a mean sequencing depth of ∼150,000 reads per cell, we detected an average of 18,661 UMIs and 6,236 genes per cell (Fig. 3b,c).

**Fig. 3.**
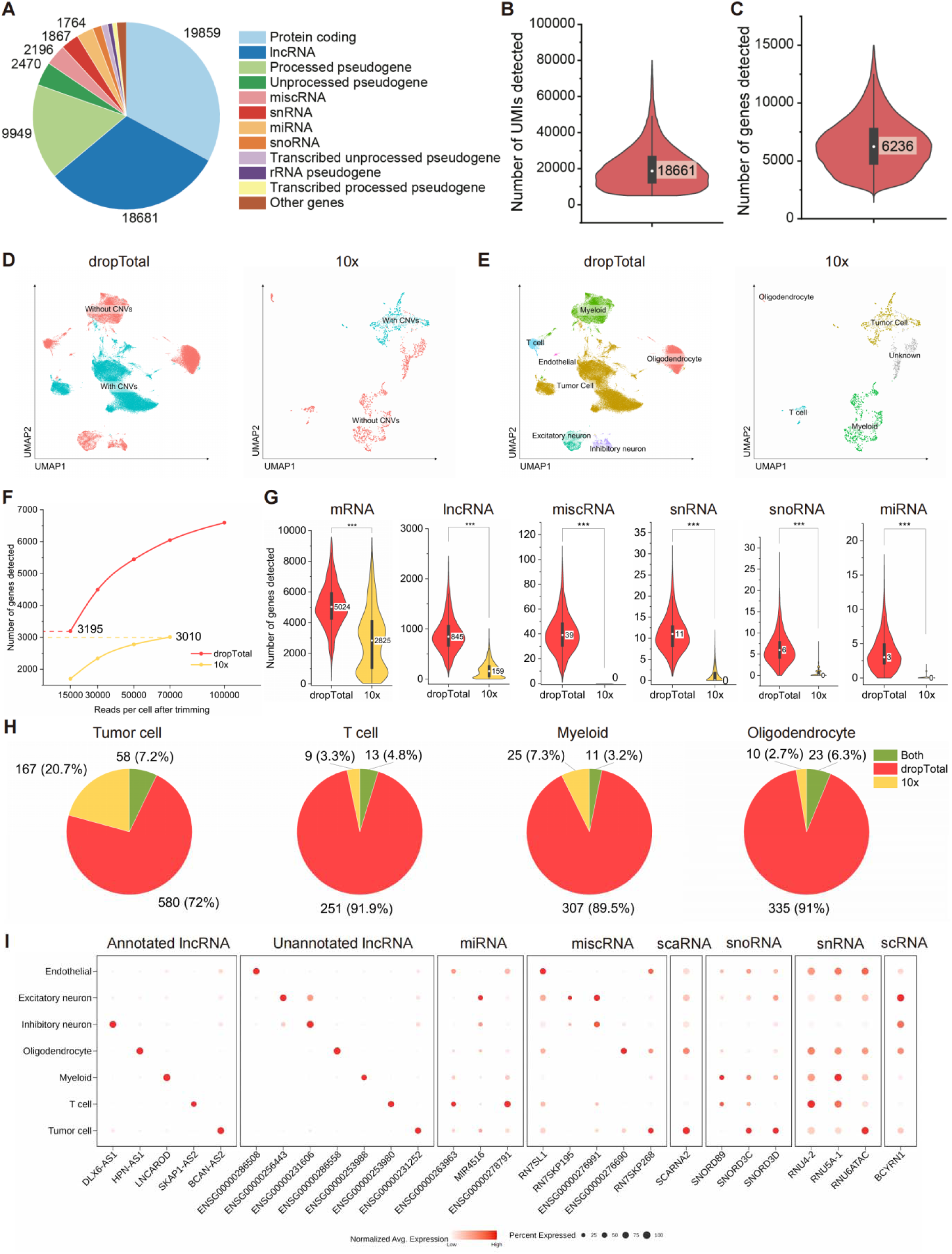
Performance of dropTotal on frozen clinical samples and comparison with 10x Genomics. (A) Proportion of detected RNA categories in gliomas. (B and C) Detection sensitivity of gliomas in UMI counts (B) and gene counts (C). (D) UMAP visualization of CNVs inferred by InferCNV (left: dropTotal; right: 10x). (E) UMAP visualization of cell types in gliomas. (left: dropTotal; right: 10x). (F) Comparison of downsampling-based gene detection sensitivity. (G) Number of mRNA, lncRNA, miscRNA, snRNA, snoRNA and miRNA genes detected by dropTotal and 10X at depth of 70k reads. (H) Cell-type differentially expressed non-coding genes identified by dropTotal and 10x. (I) Cell-type-specific expression of non-coding RNAs. Statistics test: two sample t-test in (G), with ***p<0.0001.

To further assess the performance of dropTotal on frozen clinical samples, we analyzed single-cell data from 10x Genomics-processed glioma samples using an identical workflow for direct comparison. Leveraging the hallmark chromosomal alterations of GBM—gain of chromosome 7 and loss of chromosome 10—we distinguished tumor cells from normal cells based on inferred copy-number variations (CNVs) derived from inferCNV^17,18^ (Fig. 3d and Extended Data Fig. 2). Cell-type annotation using coding-gene markers revealed that all cell populations identified by dropTotal, with the exception of endothelial and neuronal cells, were concordant with those identified by the 10x, demonstrating strong agreement between dropTotal and the current gold-standard method in the resolution of major cell types (Fig. 3e and Extended Data Fig. 3).

Given the discrepancy in sequencing depth between the two datasets, we performed downsampling analysis to enable a normalized comparison of sensitivity. Notably, dropTotal achieved detection of 3,195 genes at a depth of 15,000 reads per cell, already exceeding the 3,010 genes detected by the 10x platform at 70,000 reads per cell, indicating that dropTotal offers approximately five-fold greater sensitivity than the 10x Genomics (Fig. 3f). At the sequencing depth of 70,000 reads per cell, we compared gene detection across different RNA types and found that dropTotal exhibited significantly higher sensitivity for all classes of transcripts. Strikingly, whereas the 10x yielded a median detection count of zero for sncRNAs, dropTotal robustly detected substantial numbers of miscRNAs, snRNAs, snoRNAs, and miRNAs (Fig. 3g). Collectively, these results provide compelling evidence that dropTotal achieves high sensitivity for both coding and non-coding RNAs, with particularly pronounced advantages for non-coding transcripts, thereby addressing a critical gap in single-cell transcriptomic profiling of the non-coding transcriptome.

dropTotal consistently detects non-coding RNAs with cell-type-specific expression patterns, underscoring the potential role of non-coding transcripts in maintaining cellular identity. Using the Wilcoxon rank-sum test, we identified hundreds of differentially expressed non-coding RNAs across distinct cell populations (avg_log2FC > 0.5, min.pct = 0.25, p_val_adj < 0.05). Applying the same statistical thresholds to the 10x dataset for comparison, we observed that among three shared normal cell types, dropTotal identified approximately ten times as many differentially expressed non-coding RNAs as the 10x. In tumor cells, the number of differentially expressed non-coding RNAs captured by dropTotal was threefold higher than that detected by 10x (Fig. 3h). Representative examples include *BCAN-AS2* in tumor cells, *SKAP1-AS2* in T cells, *LNCAROD* in myeloid cells, *HPN-AS1* in oligodendrocytes, and *DLX6-AS1* in inhibitory neurons. Furthermore, we discovered numerous lncRNAs that are not yet annotated in the reference genome yet exhibit marked cell-type enrichment, such as *ENSG00000231252* in tumor cells, *ENSG00000253980* in T cells, *ENSG00000253988* in myeloid cells, *ENSG00000286558* in oligodendrocytes, *ENSG00000231606* in inhibitory neurons, *ENSG00000256443* in excitatory neurons, and *ENSG00000286508* in endothelial cells. Additionally, we identified cell-type-specific expression of sncRNAs, including *RN7SKP268* in tumor cells, *ENSG00000278791* in T cells, *RNU3A-1* in myeloid cells, *RNU4-2* in oligodendrocytes, *ENSG00000276991* in inhibitory neurons, *MIR4516* in excitatory neurons, and *RN7SL1* in endothelial cells (Fig. 3i). Collectively, these findings demonstrate that dropTotal substantially expands the repertoire of non-coding cell-type markers across all major cell populations and establishes the method as a powerful tool for probing non-coding RNA function and identifying potential therapeutic targets.

### Co-expression network reveals non-coding RNA-driven differential gene modules in primary and recurrent oligodendroglioma

To investigate the coordinated regulatory interplay between non-coding and coding RNAs, we performed weighted gene co-expression network analysis (WGCNA) on tumor cells from OG samples and identified 20 distinct gene modules (Fig. 4a). Several of these modules exhibited expression patterns specifically associated with primary versus recurrent states. Notably, M17 was markedly upregulated in primary tumors, whereas M5, M8, and M14 displayed pronounced upregulation in recurrent tumors. In addition, modules M6 and M13 exhibited strong co-expression correlation and consistent expression patterns (Fig. 4b,c and Extended Data Fig. 4).

**Fig. 4.**
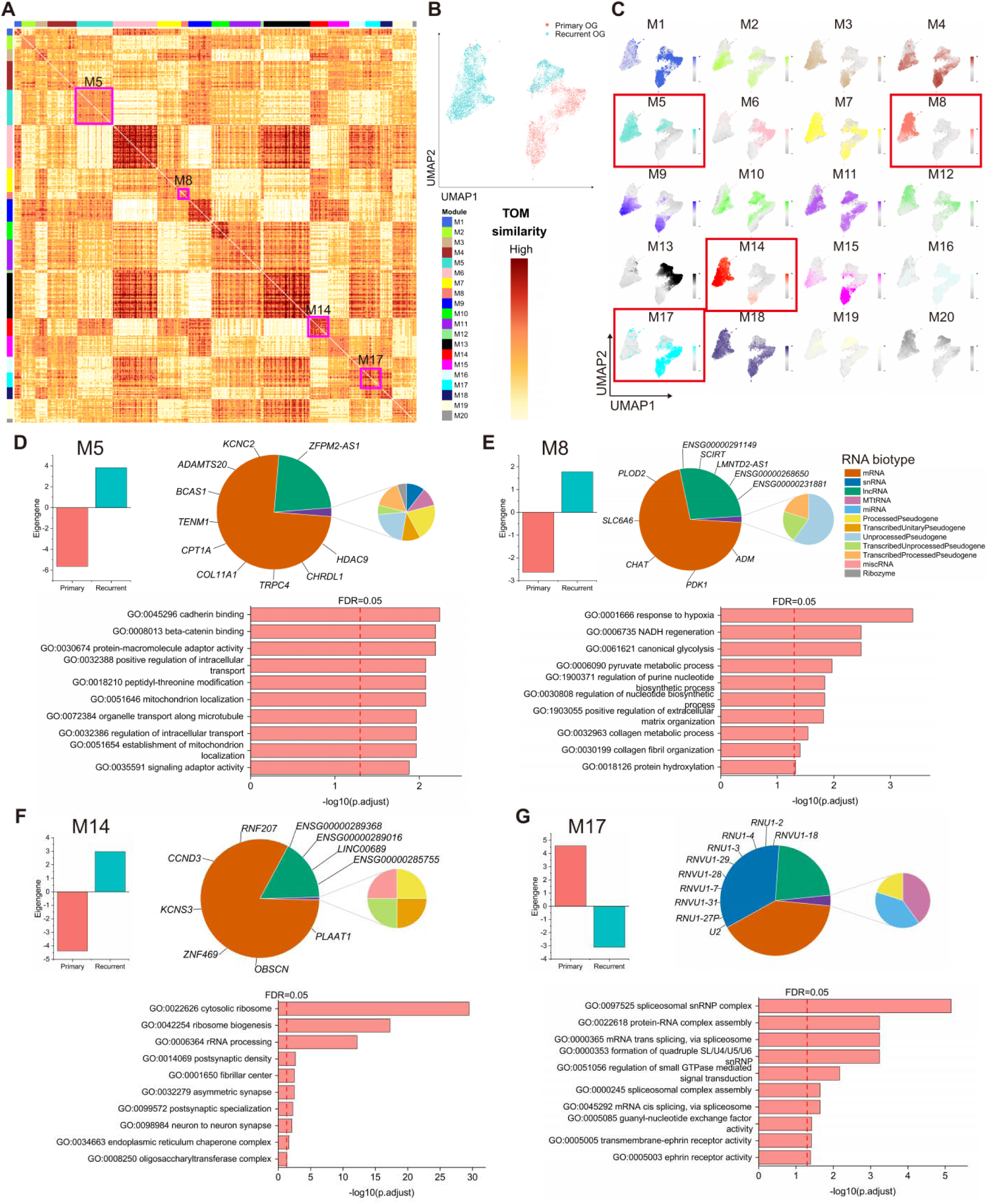
Co-expression analysis of primary and recurrent oligodendroglioma. (A) TOM heatmap of oligodendroglioma tumor cells. (B) UMAP visualization of primary and recurrent oligodendroglioma tumor cells. (C) UMAP visualization of gene module eigengene score. (D-G) Gene co-expression module 5 (D), module 8 (E), module 14 (F) and module 17 (G). Top left: module eigengene score in primary and recurrent oligodendroglioma. Top right: RNA biotype composition of the module. Top 10 hub genes of module are shown. Bottom: GO enrichment terms for genes in the module, assessed using a one-sided hypergeometric test with P values adjusted for multiple comparisons using the Benjamini–Hochberg FDR < 0.05.

Further examination of the four modules that were significantly upregulated in either primary or recurrent OG revealed that each module comprised both non-coding and coding genes. We extracted the top ten hub genes ranked by intramodular connectivity for each module and found that all modules contained ncRNAs among their hub genes (Fig. 4d-g). Strikingly, in the M17, all top ten hub genes were snRNAs (Fig. 4g). snRNAs are essential constituents of the spliceosome, typically lack poly(A) tails, and range from only 100 to 200 nucleotides in length. As a result, they are almost entirely missed by conventional single-cell transcriptomic methods such as the 10x. The efficient capture of these snRNAs by dropTotal enabled their identification as central network hubs, thereby uncovering a critical regulatory role for snRNAs in primary OG.

Gene Ontology (GO) enrichment analysis of the module-specific gene sets revealed the following associations. In M5, the hub ncRNA included *ZFPM2-AS1*, and the module was enriched for pathways related to tumor microtube network formation, β-catenin signaling activation, and mitochondrial spatial repositioning (Fig. 4d). In M8, the hub ncRNAs comprised *ENSG00000291149*, *SCIRT*, *LMNTD2-AS1*, *ENSG00000268650* and *ENSG00000231881*; this module was enriched for hypoxia response, aberrant extracellular matrix remodeling, and increased DNA repair demand (Fig. 4e). M14 contained hub ncRNAs *ENSG00000289368*, *ENSG00000289016*, *LINC00689*, and *ENSG00000285755*, with enrichment for ribosome biogenesis and translational reprogramming, tumor–neuron synaptic connectivity, and enhanced endoplasmic reticulum quality control and stress resilience (Fig. 4f). M17 harbored hub ncRNAs including *RNVU1-18*, *RNU1-2*, *RNU1-4*, *RNU1-3*, *RNVU1-29*, *RNVU1-28*, *RNVU1-7*, *RNVU1-31*, *RNU1-27P*, and *U2*; this module was enriched for RNA splicing dysregulation and spliceosome assembly, Ephrin receptor signaling, and small GTPase-mediated signal transduction (Fig. 4g). Integrating these pathway enrichment results with prior studies, we hypothesize that in primary OG, tumor cells exploit global epigenetic dysregulation—triggered by events such as IDH mutation—to extensively remodel the alternative splicing landscape, generating aberrant protein isoforms conducive to survival, while simultaneously reactivating embryonic Ephrin guidance cues to facilitate diffuse infiltration along white matter tracts^19–21^. In recurrent tumors, by contrast, cells adopt a distinct survival strategy: in response to hypoxia resulting from vascular compromise, they enhance glycolytic metabolism and secrete collagenous barriers to resist chemotherapeutic agents and immune attack; they also extend tumor microtubes that form synaptic connections with neurons, hijacking normal neural signaling to drive proliferation; and to sustain this highly invasive and recurrent state, ribosomes operate at elevated capacity for protein synthesis while nucleotide biosynthesis is accelerated to repair DNA damage^22–25^.

We also observed that M6 and M13 displayed highly similar expression patterns and strong inter-module connectivity across a subset of both primary and recurrent tumor cells (Fig. 4b and Extended Data Fig. 4). Hub ncRNAs in M6 included *LINC02058* and *LINC01151*, and the module was predominantly enriched for pathways related to angiogenesis and collagen-containing extracellular matrix organization. M13 was enriched for pathways governing small GTPase/Rho/Ras signaling and vascular permeability regulation (Extended Data Fig. 5). The coordinated co-expression of these two modules indicates that OG cells—irrespective of disease stage—are fundamentally dependent on perivascular niche remodeling and vessel co-option for widespread parenchymal dissemination. Tumor cells employ Rho/Ras family small GTPase signaling to drive cytoskeletal rearrangement and focal adhesion assembly, thereby acquiring the motile capacity for rapid migration along the abluminal surfaces of blood vessels^26,27^. Concurrently, the synergistic action of these modules actively disrupts blood–brain barrier integrity, facilitating nutrient acquisition and creating a microenvironment permissive to tumor invasion^28^.

Collectively, these findings demonstrate that ncRNAs are systematically co-expressed with coding genes and play essential functional roles in both primary and recurrent OG. This further substantiates that dropTotal, by virtue of its efficient capture of both coding and non-coding RNAs, enables sensitive detection and mechanistic elucidation of ncRNA function directly from clinical samples.

### dropTotal dissects the differential regulatory functions of non-coding RNAs across six cellular states of glioblastoma

To investigate the regulatory roles of ncRNAs in GBM tumor cells, we first annotated six distinct cellular states—MES-like 1, MES-like 2, AC-like, NPC-like 1, NPC-like 2, and OPC-like—based on established marker gene signatures^29^. Using the Wilcoxon rank-sum test, we identified hundreds of ncRNAs that were differentially expressed across these cellular states (avg_log FC > 0.5, min.pct = 0.25, p_val_adj < 0.05) (Fig. 5a, Extended Data Fig. 6 and Extended Data Fig. 7). Representative examples of annotated non-coding RNAs with state-specific enrichment include *MIR222HG* in MES-like 1; *ADAMTS9-AS2* and *SCIRT* in MES-like 2; *LINC00595* in AC-like; *MACORIS* in NPC-like 1; *DLX6-AS1* and *KCNH7-AS1* in NPC-like 2; and *MIR219A2HG* in OPC-like cells. In addition, we detected a substantial number of ncRNAs that are currently absent from reference genome annotations, including *ENSG00000255029* in MES-like 1, *ENSG00000273184* in MES-like 2, *ENSG00000286757* in AC-like, *ENSG00000231918* NPC-like 1, *ENSG00000236501* in NPC-like 2, and *ENSG00000266844* in OPC-like cells. Notably, the majority of the top-ranked differentially expressed ncRNAs in each cellular state corresponded to unannotated transcripts (Fig. 5b).

**Fig. 5.**
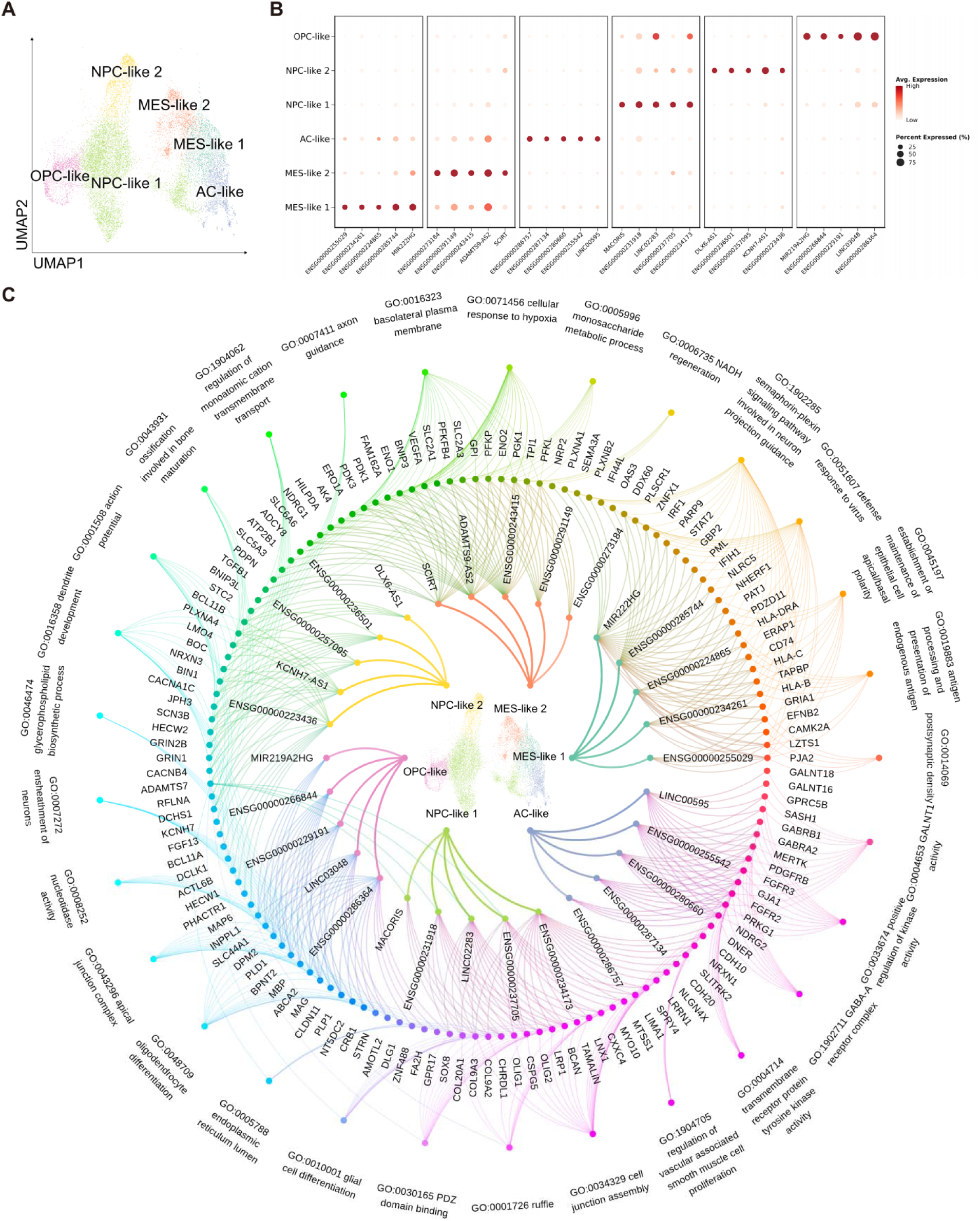
Predicted functions of cellular-state-specific non-coding RNAs. (A) UMAP visualization of six cellular states in glioblastoma tumor cells. (B) Cellular-state-specific non-coding RNA expression. (C) Functional identification of cellular-state-specific non-coding RNAs. From inner to outer: cellular state, cellular-state-specific non-coding RNAs, target mRNAs, and GO enrichment pathways.

Having identified state-specific ncRNAs, we next sought to infer their potential regulatory functions within each cellular state. For each cellular state, we used GRNBoost2 to infer weighted gene–gene associations between selected ncRNAs and other expressed genes. Edges were ranked according to GRNBoost2 feature-importance weights, and the top 50 associated genes for each ncRNA were retained for GO enrichment analysis. (Fig. 5c).

Within the MES-like states, core ncRNAs of the MES-like 1 subpopulation—such as *ENSG00000234261*, *ENSG00000285744*, and *MIR222HG*—predominantly targeted *HLA* family genes, *STAT2*, and *IFIH1*, exhibiting specific enrichment for antigen presentation and antiviral defense response pathways, thereby mediating tumor–immune crosstalk. In contrast, non-coding RNAs characteristic of the MES-like 2 subpopulation—including *SCIRT*, *ADAMTS9-AS2*, *ENSG00000291149*, and *ENSG00000243415*—selectively targeted genes such as *VEGFA*, *PGK1*, and *PFKP*, with enrichment in cellular response to hypoxia and monosaccharide metabolic processes, driving metabolic reprogramming to accommodate the hypoxic microenvironment^30^. Analogous network heterogeneity was observed among progenitor-like states. The OPC-like state was primarily governed by *ENSG00000266844*, *ENSG00000286364*, and *LINC03048*, which preferentially targeted *MBP*, *MAG*, and *PLP1* to sustain myelination programs and glial differentiation characteristics. The NPC-like states exhibited further functional divergence: ncRNAs in NPC-like 1 (e.g., *MACORIS*, *LINC02283*) tended to regulate *ADAMTS7* and *OLIG1*, participating in glial differentiation and cell junction assembly, whereas those in NPC-like 2 (e.g., *DLX6-AS1*, *KCNH7-AS1*, and *ENSG00000223436*) specifically targeted *KCNH7*, *SCN3B*, and *PLXNA4*, with prominent enrichment in action potential propagation, axon guidance, and dendrite development—thereby providing molecular infrastructure for the construction of pro-tumorigenic electrophysiological networks^22^. Furthermore, the AC-like state was dominated by *ENSG00000287134* and *ENSG00000255542*, which converged on key genes such as *FGFR3*, *PDGFRB*, and *MERTK* to maintain transmembrane receptor protein tyrosine kinase activity and sustain canonical proliferative and invasive signaling.

Collectively, these analyses demonstrate that a repertoire of state-specific ncRNAs in GBM orchestrates highly modular “ncRNA–target gene–functional pathway” axes, thereby heterogeneously shaping and perpetuating the core biological properties of distinct tumor cell subpopulations.

### Cellular-state-specific alternative splicing landscape mediated by non-coding RNAs in glioblastoma

As an essential post transcriptional gene regulation mechanism, Alternative Splicing (AS) not only serves as an essential process regulating neurogenesis^31–34^, but also plays a significant role in tumorigenesis, affecting tumor cell proliferation, apoptosis, invasion, metastasis, etc^35^. The ability to profile full transcripts allowed dropTotal to capture AS events and identify their patterns and potential regulatory splicing factors (SFs) across different cellular status in GBM tumor cells. To overcome the limited coverage of splicing nodes at single-cell level, a random pooling pseudo-bulk approach from MicroExonator was applied^36^. Single cells from the same cell type were randomly selected and their reads were pooled for percent spliced in (PSI) quantification using Whippet. To avoid false positives due to pooling, the process was repeated 50 times.

To reveal general splicing patterns across 6 cellular states in tumor cells derived from GBM, we identified splicing nodes showing significant deviation of PSI values from the rest of cell types, which are denoted as splice node markers (SNMs). We totally identified 428 SNMs (226 specifically included nodes and 202 specifically excluded nodes, Fig. 6a,b). NPC-like-2 state contributes 40% of total SNMs detected (172 SNMs), exhibiting extensive AS activity. It has previously been shown that GBM recapitulates a normal neurodevelopment trajectory^37^. Moreover, AS has long been recognized as a key regulatory mechanism in neurogenesis and brain development, especially during NPC differentiation. NPC-like-2 state potentially recapitulated this process in GBM.

**Fig. 6.**
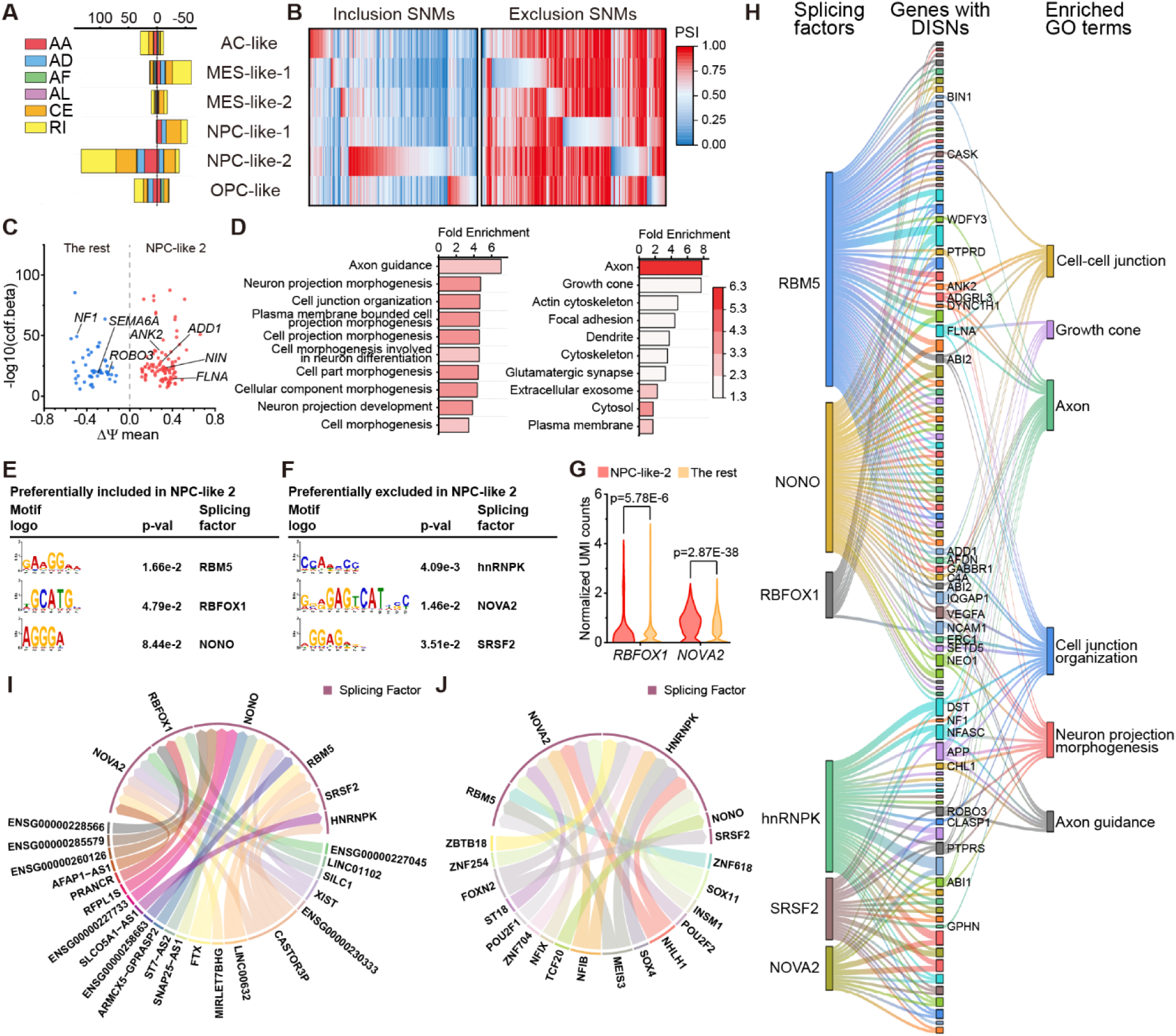
Alternative-splicing events and predicted splicing regulation in glioblastoma. (A) Number of SNMs identified in 6 cellular states of GBM tumor cells. (B) Heatmap showing ψ values across cellular states corresponding to (A). (C) Volcano plot illustrating DISNs identified between NPC-like-2 and the rest states. (D) GO enrichments of genes with DISNs identified between NPC-like-2 and the rest states using biological processes (left) and cellular compartments (right). (E) Top 3 splicing factor binding motifs enriched in 200 bp upstream of differentially included nodes in NPC-like-2 state. (F) Top 3 splicing factor binding motifs enriched in 200 bp downstream of differentially excluded nodes in NPC-like-2 state. (G) Violin plot illustrating significant upregulation of RBFOX1 and NOVA2 in NPC-like-2 against the rest states. (H) Sankey plot illustrating composition of splicing factor binding motifs enriched in 200 bp upstream of differentially included nodes and 200 bp downstream of differentially excluded nodes in NPC-like-2, as well as GO enrichment of genes with DISNs. (I) Top-ranked regulatory atlas of lncRNAs against SFs in NPC-like-2. (J) Top-ranked regulatory atlas of TFs against SFs in NPC-like-2.

Next, we identified differentially included splicing nodes (DISNs) in 15 comparisons among 6 cellular states in GBM cells. The number of DISNs identified ranged from 492 to 1773, indicating AS occurs frequently in GBM (Extended Data Fig. 8). To gain more biological insights into those DISNs, we performed GO enrichment analysis on all 15 comparisons. For the majority of these comparisons, terms related to normal neurogenesis were enriched (Extended Data Fig. 9), which is consistent with the previous finding that GBM partially mimicked the neuronal development. Interestingly, we observed numerous cell migration related terms as well as EMT transition enriched in the comparison between MES-like-1 and MES-like-2 state (Extended Data Fig. 8c,d). This agrees with our previous finding that MES-like state is active for EMT transition, showing that EMT pathway is also regulated through AS, which was captured by our method.

Since NPC-like-2 state exhibited the most abundant AS events, we seek to construct the regulatory atlas of AS in NPC-like-2 state. We first identified 154 DISNs in NPC-like-2 against all other states. From all genes affected by those DISNs we noticed multiple genes that are important for neuronal development: *ADD1*, *ANK2*, *NIN*, *FLNA*, *ROBO3*, *NF1* and *SEMA6A*, most of which are known to harbor AS events^31,38–40^. GO analysis also revealed the genes affected by the DISNs were enriched in axon for cellular compartments and axon guidance for biological pathways (Fig. 6d). To identify potential regulatory SFs, sequences of 200 bp upstream and downstream of splicing nodes were extracted and analyzed by SEA from MEME suite. *RBM5*, *RBFOX1* and *NONO* were enriched in upstream of preferentially included nodes while *HNRNPK*, *NOVA2* and *SRSF2* were enriched in downstream of preferentially excluded nodes (Fig. 6e,f), among which *RBFOX1* and *NOVA2* were upregulated in NPC-like-2 (Fig. 6g). *RBFOX1*, *NONO*, *HNRNPK* and *NOVA2* were previously known to regulate neuronal development process through splicing^41–44^. To examine the biological functions of the nodes potentially regulated by these SFs, we annotated the nodes with regulatory SFs and GO terms on a Sankey plot (Fig. 6h). To elucidate the regulators of SFs, we constructed regulatory atlas of SF as previously described (Fig. 6I,j). Among the potential regulators: (i) the murine homology of lncRNA *SILC1* was required in regenerating neuron^45^; (ii) TF *SOX4* and *SOX11* are vital for neuronal protein expression^46^, while *POU2F1* and *POU2F2* are also essential for neuronal lineage differentiation^47,48^. Taken together, we show that profiling full length transcripts with high sensitivity using dropTotal enabled identification of general AS patterns and DISNs across different cellular states in GBM, as well as revealing potential regulatory splicing factors.

## Discussion

Here, we introduce dropTotal, a high-throughput droplet-based single-cell total RNA-sequencing method that couples temperature-ramped hybridization of a dU-modified GAT primer with physically separated reverse-transcription and barcoding reactions. In benchmarked cell lines, dropTotal detected more genes per cell than the other methods evaluated, while providing broad gene-body coverage and substantially improving recovery of lncRNAs and sncRNAs. These results show that efficient capture of both polyadenylated and non-polyadenylated transcripts can extend single-cell analysis beyond conventional mRNA-centered profiles.

Application to frozen human glioma nuclei illustrates the biological value of this increased sensitivity. dropTotal recovered the major cell populations detected by a 10x Genomics workflow but identified a much broader repertoire of cell-type-enriched ncRNAs, including short and incompletely annotated transcripts. Thus, the principal advantage is not simply a larger feature count: improved ncRNA detection exposes molecular variation that can be organized by cell type, disease stage and tumor cell state.

In oligodendroglioma, co-expression analysis identified modules associated with primary or recurrent tumors and revealed ncRNAs among their most highly connected genes, including a primary-tumor module dominated by snRNAs. In glioblastoma, state-enriched ncRNAs and GRNBoost2-derived gene-association weights suggested distinct functional programs across six tumor cell states, spanning immune response, hypoxia adaptation, lineage identity and neuronal-like signaling. These network results nominate testable ncRNA–gene relationships rather than establishing direct regulation or causality.

The broad gene-body coverage of dropTotal also enabled analysis of alternative splicing. The state-specific splice-junction markers, particularly those enriched in the NPC-like 2 state, connected neuronal-development and axon-guidance programs with candidate splicing factors and upstream regulators. This analysis illustrates how ncRNA expression and isoform-level variation can be examined in the same single-cell dataset, although the inferred regulatory links require experimental validation.

Several limitations remain. The glioma analyses were performed in a limited clinical cohort and should be validated in larger, independent datasets. Comparisons with published methods may also retain platform-and dataset-specific effects despite downsampling. Network inference, motif enrichment and pseudo-bulk splicing analyses are associative, and perturbation experiments will be needed to determine which ncRNAs are functional regulators. In addition, the performance of dropTotal across fresh, fixed and FFPE material, and its integration with spatial or epigenomic measurements, warrants systematic benchmarking.

Overall, dropTotal converts efficient ncRNA detection into cell-type-resolved atlases, candidate regulatory networks and cell-state-resolved splicing maps. Its application should facilitate systematic investigation of coding and non-coding transcript regulation in heterogeneous tissues.

## Methods

### Cell culture

HEK293T and NIH/3T3 cells were cultured in high-glucose DMEM supplemented with 10% fetal bovine serum and 1× penicillin–streptomycin. Cells were passaged every 2–3 days.

### Human glioma samples

A total of 26 human glioma specimens were obtained from 13 adult participants who underwent paired surgical resections for primary and recurrent glioma at Shandong Provincial Hospital Affiliated to Shandong First Medical University. Each participant contributed one primary and one recurrent tumor specimen. The cohort comprised nine male and four female participants. Ages at the primary resection ranged from 38 to 66 years (median, 50 years), and ages at the recurrent resection ranged from 41 to 67 years (median, 53 years). Diagnoses included astrocytoma, glioma, oligodendroglioma with 1p/19q codeletion and glioblastoma, encompassing CNS WHO grades 2–4. Detailed demographic and clinicopathological characteristics of each specimen are provided in Supplementary Table 1.

Primary specimens were defined as tissues collected during the initial surgical resection, whereas recurrent specimens were collected during a subsequent resection following tumor recurrence. The collection and use of human samples were approved by the Institutional Review Board of Shandong Provincial Hospital Affiliated to Shandong First Medical University (NSFC NO: 2024–016).

### Single nuclei preparation from frozen human glioma samples

Nuclei were isolated from frozen brain samples using the protocol of Krishnaswami et al^49^. Briefly, tissue was homogenized in homogenization buffer with a Dounce homogenizer and passed through a 40-µm filter. Nuclei were fixed with 3% paraformaldehyde for 10 min at room temperature and quenched with 2.5 M glycine. After washing, nuclei were stained with Hoechst 33342 and enriched by fluorescence-activated cell sorting on a Beckman Coulter CytoFLEX SRT. Nuclei were resuspended in PBS containing 9% Ficoll 400, counted and diluted to 1.6 × 10^6 nuclei ml−1 for droplet generation.

### Microfluidic device design and fabrication

The microfluidic channel layout was designed using AutoCAD and printed on a high-resolution photomask. Microfluidic molds were patterned in SU-8 3050 to a height of 80 µm according to the manufacturer’s instructions. PDMS devices were prepared from RTV615 silicone elastomer base and curing agent at a 10:1 (w/w) ratio and cured at 80 °C for at least 2 h. Inlets and outlets were made with a 1-mm biopsy punch. The patterned PDMS chip was plasma-bonded to a glass slide, and the channel surface was treated with trimethylchlorosilane to render it hydrophobic.

### Barcoded hydrogel beads synthesis

Hydrogel-bead and barcode synthesis followed a previously reported protocol. A deoxyuridine was incorporated into the acrydite-modified primer to permit enzymatic release of barcoded primers by USER enzyme. Two rounds of split-and-pool synthesis were performed, with 768 barcode sequences in each round, yielding 589,824 barcode combinations. In each round, hydrogel beads were distributed across 768 wells, annealed to templates containing well-specific barcodes and extended with Bst 2.0 WarmStart DNA polymerase for 3 h at 52 °C. After cleanup, barcoded hydrogel beads were transferred to bead-freezing buffer and stored at −20 °C. Before use, beads were washed twice in 1× ThermoPol buffer.

### dropTotal procedures

#### Cell encapsulation and reverse transcription

Dissociated cells were washed twice with PBS, counted, and adjusted to 1.6 × 10^6^ cells ml^−1^. For the species-mixing experiment, HEK293T and NIH/3T3 cells were combined at a 1:1 ratio. To minimize sedimentation during encapsulation, the cell suspension was mixed with OptiPrep before loading.

Cells and reverse-transcription (RT) reagents were co-encapsulated in droplets using a microfluidic device with HFE7500 oil containing 2% fluorosurfactant (Supplementary Video 1). The collected droplets were subjected to a multiple-annealing RT program, followed by heat inactivation.

#### Single-cell cDNA barcoding

Prepared barcoded hydrogel beads (BHBs) and the barcoding reaction mixture were loaded into a droplet-merging microfluidic device. Droplets containing individual BHBs were paired and merged with RT droplets, after which the merged droplets were stabilized with fluorosurfactant-containing oil (Supplementary Video 2).

Within the merged droplets, barcoded primers were released from the BHBs, unused RT primers were degraded by USER enzyme, and RNA templates were removed using RNase H and RNase If. Barcoded primers were subsequently annealed to and extended along the single-stranded cDNA templates.

#### Post-barcoding processing

Following barcoding, droplets were demulsified using perfluoro-1-octanol in TE buffer under conditions designed to rapidly terminate polymerase activity and minimize barcode crosstalk. BHBs were removed by centrifugation, and the recovered cDNA was purified using DNA selection beads. Residual barcoded primers were blocked by terminal transferase-mediated incorporation of ddCTP, followed by bead-based purification.

#### Library construction and sequencing

Sequencing libraries were constructed by adding the Read 2 sequencing handle to the barcoded cDNA, followed by pre-amplification and bead-based purification. Illumina-compatible i5 and i7 indices were subsequently introduced by PCR. Library yield was determined using the Qubit dsDNA HS assay.

Final libraries were pooled and sequenced on the GeneMind SURFSeq 5000 platform using paired-end 150-bp reads.

### dropTotal raw data pre-processing

Cutadapt (version 3.4) was used to trim the residual RT primer sequence from the 5’ of Read 2 in a paired-read mode. UMI-tools (version 1.1.2) ‘whitelist’ command was used to determine a cell barcode combination list. Potential errors introduced during barcoded primer synthesis and library construction, a custom Python script was used to correct the cell barcode list by comparing to all known cell barcodes. The corrected cell barcode list was provided to UMI-tools ‘extract’ command for cell barcode and UMI extraction. For Read 2, sequences matched with at least 10 nt from reverse complementary sequence of RT primer was trimmed using 3’ mode with cutadapt. In silico ribosomal depletion was performed as previously described. The processed Read 2 was mapped to the hg38 genome (or a combined genome of hg38 and mm10) using STAR (version 2.7.8a) with default parameters. The aligned reads were assigned to gene annotations (GENCODE v44 for human glioma samples, and a combined annotation of GENCODE v44 and vM33 for cell line mixing samples) using a custom python script, while giving gene annotations that do not have annotated introns higher priority. Only alignments overlapped to the gene annotations on the different strand. We found indels might prohibit reads to be aligned appropriately by STAR. Thus, we aligned the reads without any assigned gene annotations with mem algorithm in BWA (version 0.7.17). These aligned reads were assigned to gene annotations the same as described before. Reads with gene annotations assigned were combined, and the expression matrix was created with ‘count’ command from UMI-tools.

### Benchmarking against other methods

Barcodes contained more than 80% of the UMIs assigned to only one of either human or mouse were considered singlets. Barcodes with UMIs less than 20,000 were considered as backgrounds. To calculate gene body coverage, ‘geneBody_coverage.py’ from RSeQC (version 5.0.1) was used. Gene body coverage was calculated against housekeeping genes. For Smart-seq3 both reads with and without UMIs were used. To perform downsampling, reads from different barcodes were first demultiplexed into separate FASTQ files, in silico ribosomal depletion was performed and Seqtk (version 1.4) was utilized for downsampling on those FASTQ files. Single cells without sufficient reads for downsampling were discarded for the downstream analyses. To summarize different biotypes identified, expression matrix containing all single cells from either cell line or human glioma samples were used.

For the 10X dataset, Cell Ranger (version 7.2.0) was used with default parameters to perform barcode processing, mapping and expression matrix generation. Genome and gene annotation file version were kept the same as other analysis pipelines.

### scRNA-seq data analysis

#### Normalization and unsupervised clustering analysis

Gene-expression analysis was performed with Seurat v5.1.0^50^. Seurat objects were generated from the count matrices, and genes expressed in more than three cells were retained. Cells with 5,000–80,000 detected UMIs were retained. Count matrices were log-normalized with NormalizeData (scale.factor = 10,000), the 2,000 most variable genes were selected with FindVariableFeatures and the resulting matrices were scaled with ScaleData. Datasets were integrated with Harmony v1.2.0^51^ in R. Principal-component analysis was performed with RunPCA, and the number of components used downstream was selected from the elbow plot. A k-nearest-neighbor graph was constructed with FindNeighbors, clusters were identified with FindClusters (resolution = 1.0), and cells were embedded in two dimensions with RunUMAP.

#### Copy number variation (CNV) analysis

Copy-number variation (CNV) was inferred for each cell with inferCNV v1.18.1 using cutoff = 0.1, cluster_by_groups = TRUE, denoise = TRUE and HMM = TRUE. Cells with chromosome 7 gain and chromosome 10 loss in glioblastoma, or 1p/19q co-deletion in oligodendroglioma, were classified as malignant.

#### Differential expression gene analysis and cell type annotation

Differentially expressed genes defining each cluster were identified with FindAllMarkers using min.pct = 0.25 and logfc.threshold = 0.5. Genes with Benjamini–Hochberg-adjusted P < 0.05 were considered significant. Clusters were annotated from canonical cell-type markers together with inferred CNV profiles for malignant-cell identification.

#### GO enrichment analysis

The clusterProfiler (version 4.10.1) R package was applied to obtain the enriched GO terms^52^. The enriched terms were filtered by setting pvalueCutoff = 0.05, qvalueCutoff = 0.05, and were then visualized by ggplot2 (version 3.5.1) R package.

#### Subpopulation analysis of GBM cells

After cell-type annotation, tumor cells from patients with glioblastoma were selected using the pathological diagnosis. The analysis workflow described above was repeated: dimensionality reduction with RunPCA, neighborhood construction with FindNeighbors, clustering with FindClusters (resolution = 1) and marker identification with FindAllMarkers. Glioblastoma cellular states were assigned from established marker-gene expression patterns.

### hdWGCNA analysis of OG cells

hdWGCNA (high-dimensional weighted gene co-expression network analysis) is an R package designed for co-expression network analysis in high-dimensional transcriptomics data, including single-cell RNA-seq (scRNA-seq) and spatial transcriptomics. We performed hdWGCNA analysis following the standard workflow described in the official tutorial. All analyses were conducted in R environment (version 4.3.2) using the hdWGCNA package (version ≥0.2).

### Gene regulatory network inference with GRNBoost2

Analyses were performed in Python using arboreto and pySCENIC. For each glioblastoma cellular state, ncRNAs detected in more than 20% of cells were used as candidate regulators. GRNBoost2 was applied to the raw counts matrix to estimate feature-importance weights for regulator–gene relationships. Genes were ranked by importance weight for each ncRNA, and the top 50 were retained as candidate targets for GO enrichment analysis.

### Analysis of AS events across glioblastoma tumor cell subtypes

Alternative-splicing events were analyzed as previously described. Reads from randomly selected cells within each cellular state were pooled to form pseudobulk datasets, and PSI values were quantified with Whippet. Random pooling was repeated 50 times. Thresholds for identifying SNMs and DISNs followed the cited workflow. For Gene Ontology enrichment, multiple AS events assigned to the same gene were collapsed. To nominate candidate splicing factors, sequences 200 bp upstream and downstream of splice nodes were analyzed with SEA from the MEME Suite using default settings.

### Quantification and statistical analysis

Statistical analyses were performed in R v4.3.2 and Origin 2021. Differential-expression analyses used two-sided Wilcoxon rank-sum tests with Benjamini–Hochberg adjustment. Gene Ontology enrichment used one-sided hypergeometric tests with Benjamini–Hochberg correction. Cell-level comparisons in Fig. 3g used unpaired one-sided Welch’s t-tests without assuming equal variances. Test statistics and P values are provided in the relevant figure legends where available.

## Data availability

- The raw sequencing files and processed scRNA-seq data are available in BioProject database under accession no. PRJNA1402473 and are publicly available as of the date of publication. The following public datasets were used in the present study for cell line benchmark comparison: (1) VASA-drop on 293T cell line (GSE176588), (2) TotalX on 293T cell line (GSE315939), (3) 10x on 293T cell line (https://www.10xgenomics.com/datasets/500-1-1-mixture-of-human-hek-293-t-and-mouse-nih-3-t-3-cells-3-lt-v-3-1-chromium-controller-3-1-low-6-1-0), (4) MATQ-Drop on 293T cell line (GSE199346). The following public datasets were used in the present study for glioma sample benchmark comparison:10x on glioma sample (https://www.10xgenomics.com/datasets/2-k-sorted-cells-from-human-glioblastoma-multiforme-3-v-3-1-3-1-standard-6-0-0).

## Code availability

- This paper does not report original code.

## Acknowledgments

We thank the hospital for providing biological samples and patient information. We thank the group of Peiyan Ni, Associate Researcher at the Mental Health Center Affiliated to Zhejiang University School of Medicine and Hangzhou Seventh People’s Hospital, for providing the flow cytometer and assisting in the cell isolation of glioma samples. We thank Genomics Core Facility at GeneMind Biosciences Co., Ltd. (Shenzhen, China) for their assistance with sequencing. We also thank Professor Zhengdong Cheng from College of Chemical and Biological Engineering of Zhejiang University for support with computer server.

## Author Contributions

Conceptualization, X.L., W.C., Y.P., and Q.Z.; methodology, X.L., W.C., Y.P., and Q.Z.; investigation, X.L., W.C., Z.L., Y.P., T.W., Y.D. and Q.Z.; writing—original draft, X.L., W.C., and Q.Z.; writing— review & editing, X.L., W.C., and Q.Z.; funding acquisition, Y.M. and Q.Z.; resources, X.X., Z.J., C.L., Y.L., and Q.Z.; supervision, Q.Z.

## Supplementary information

Supplementary Information is available for this paper.

Supplementary Table 1. Patient clinical information related to Fig. 3.

Supplementary Video 1. Droplet generation, related to Fig. 1

Supplementary Video 2. Droplet merging, related to Fig. 1

---

**Extended Data Fig. 1.**
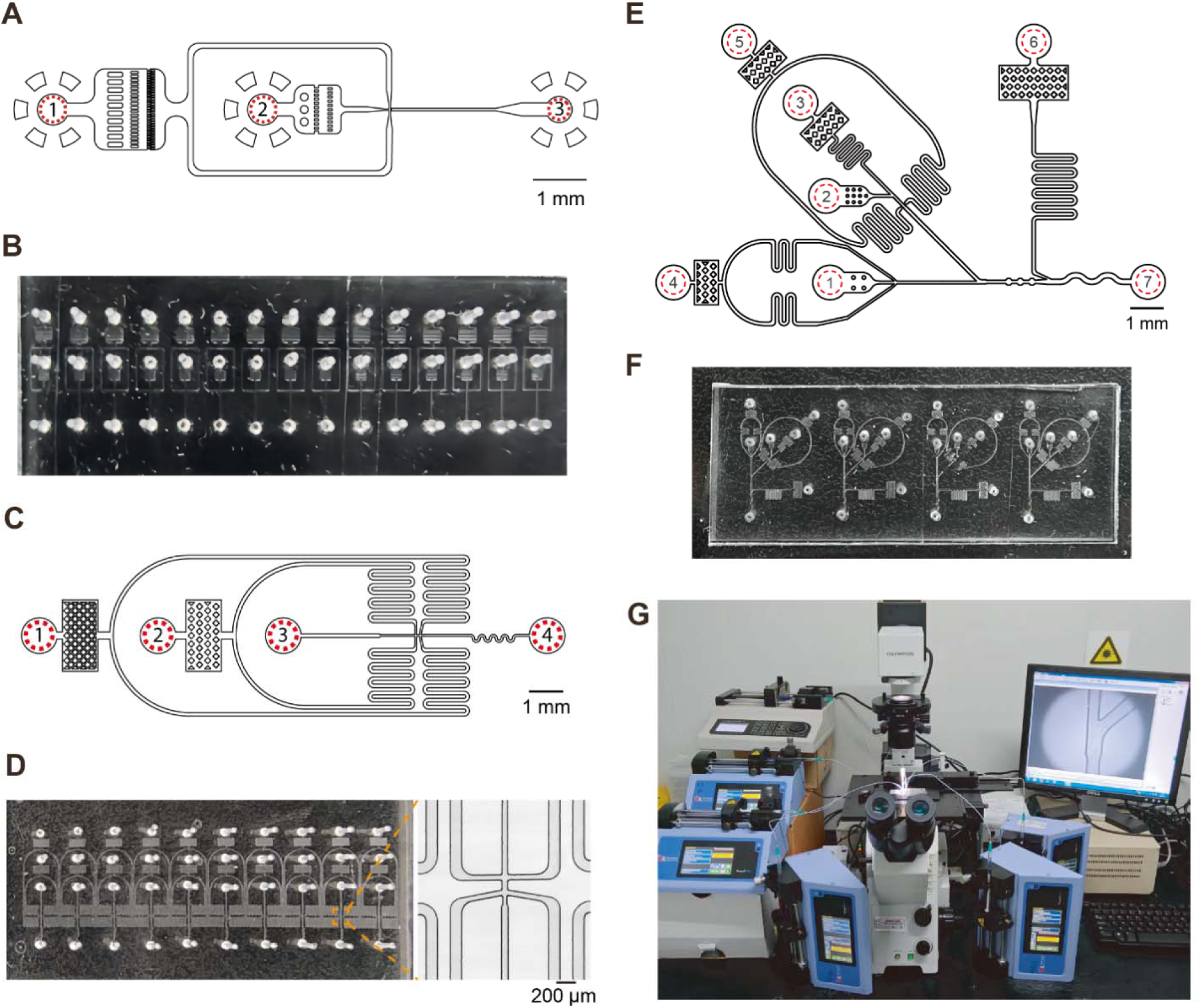
Microfluidic channel designs and dropTotal device setup. (A) Microfluidic channel design for HB droplet generation. The numbers marked indicate droplet generation oil inlet (1), acrylamide-primer mix inlet (2), and the collection outlet (3). (B) Fabricated HB droplet generation microfluidic channels. (C) Microfluidic channel design for RT droplet generation. The numbers marked indicate droplet generation oil inlet (1), RT/lysis mix inlet (2), single cell/nuclei suspension inlet (3), and the collection outlet (4). (D) Fabricated RT droplet generation microfluidic channels. (E) Microfluidic channel design for droplet merging. The numbers marked indicate RT droplet inlet (1), BHB inlet (2), barcoding reaction mix inlet (3), RT droplet separation oil inlet (4), barcoding reagent droplet generation oil inlet (5), stabilization oil inlet (6) and the collection outlet (7). (F) Fabricated droplet merging microfluidic channels. (G) Device setup for droplet merging, including 6 mechanical pumps and a microscope attached with a high-speed camera for microfluidic monitoring.

**Extended Data Fig. 2.**
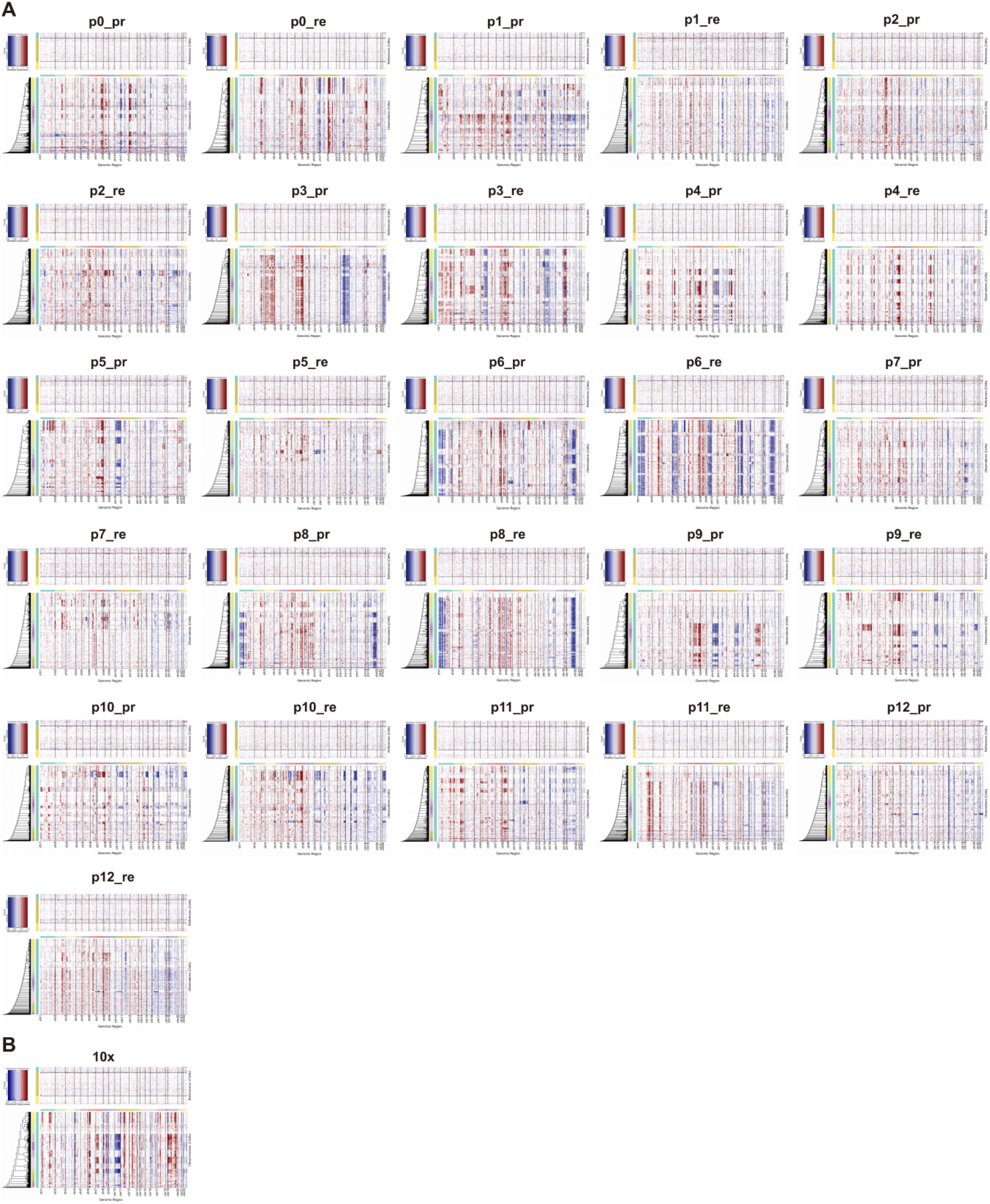
inferCNV profiles of glioma datasets generated by dropTotal and 10x Genomics.

**Extended Data Fig. 3.**
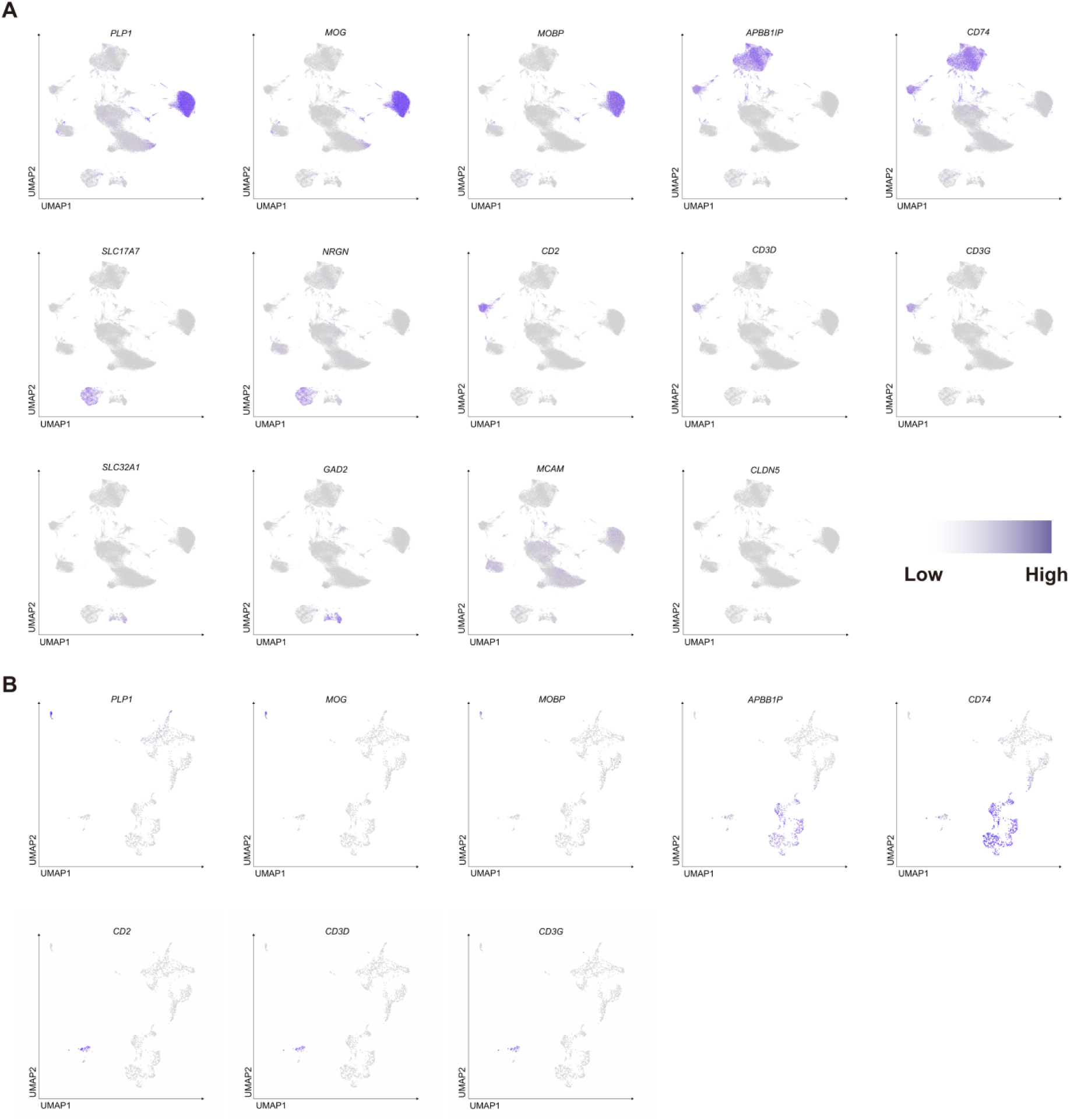
Canonical markers used to annotate normal cell types. Oligodendrocyte marker genes: *PLP1*, *MOG*, *MOBP*. Myeloid marker genes: *APBB1IP*, *CD74*. Excitatory neuron marker genes: *SLC17A7*, *NRGN*. T cell marker genes: *CD2*, *CD3D*, *CD3G*. Inhibitory neuron marker genes: *SLC32A1*, *GAD2*. Endothelial marker genes: *MCAM*, *CLDN5*.

**Extended Data Fig. 4.**
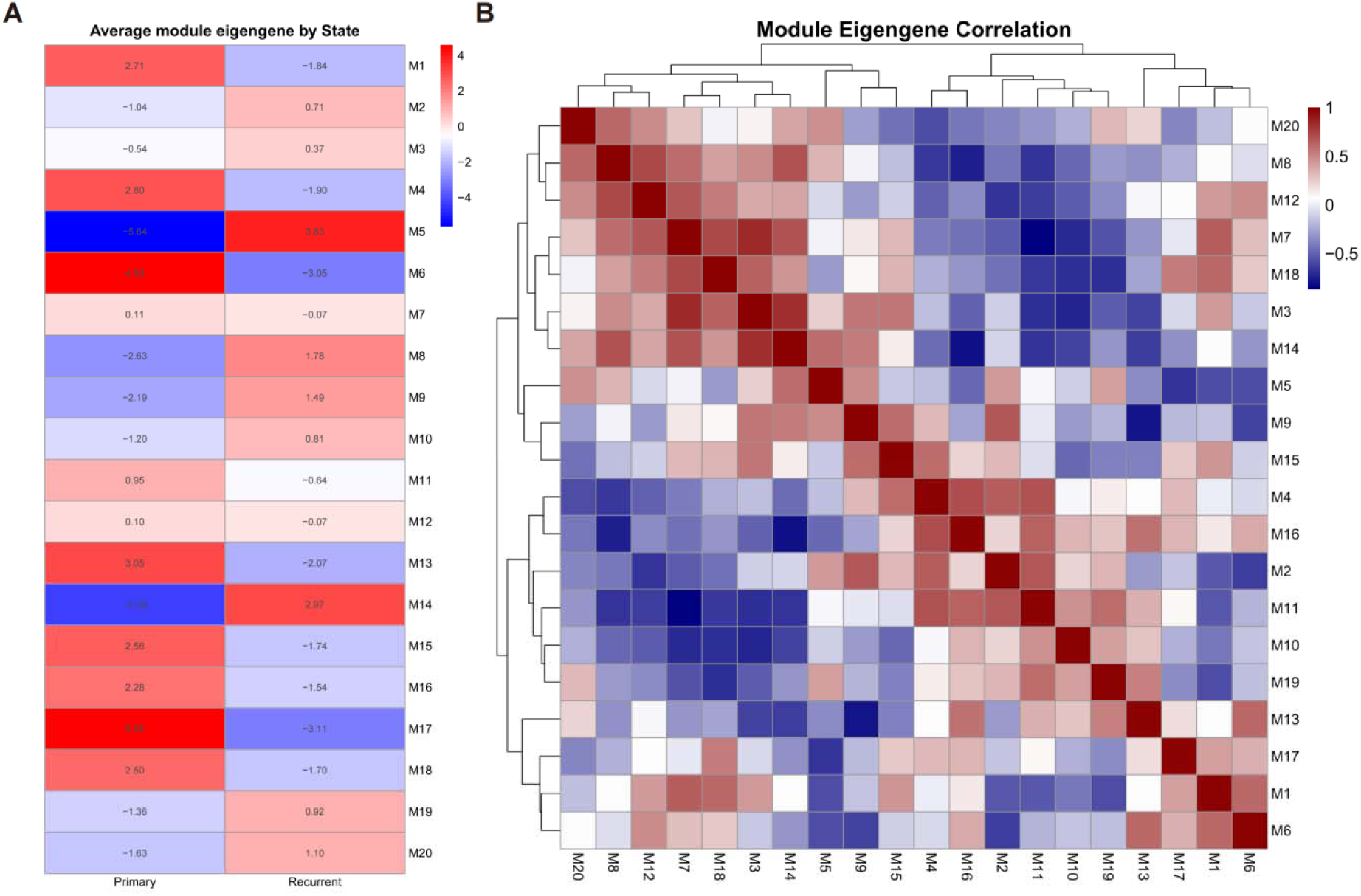
Co-expression modules in primary and recurrent oligodendroglioma. (A) Heatmap of module eigengene scores. (B) Heatmap of correlations between module eigengenes.

**Extended Data Fig. 5.**
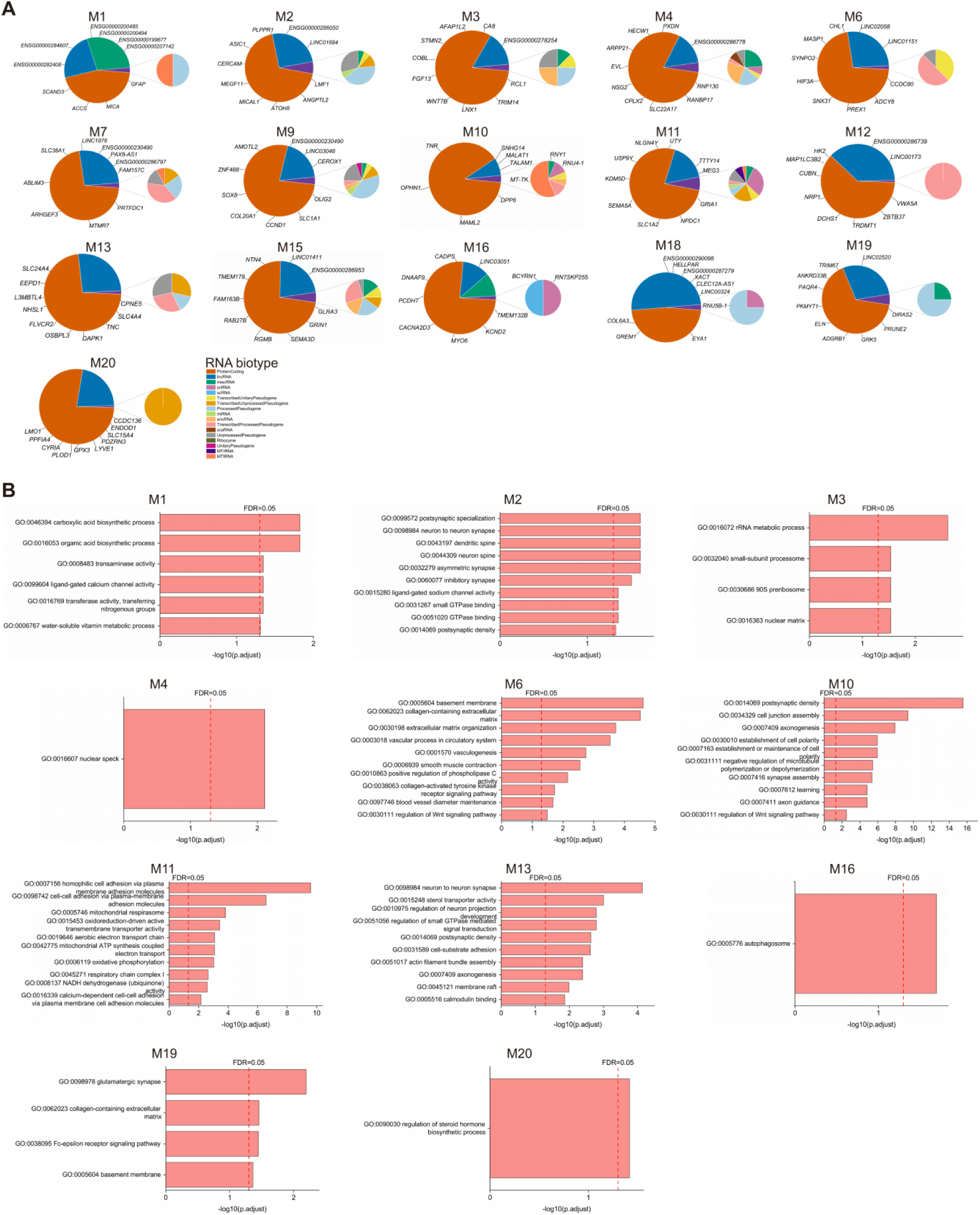
RNA composition and functional enrichment of co-expression modules.

**Extended Data Fig. 6.**
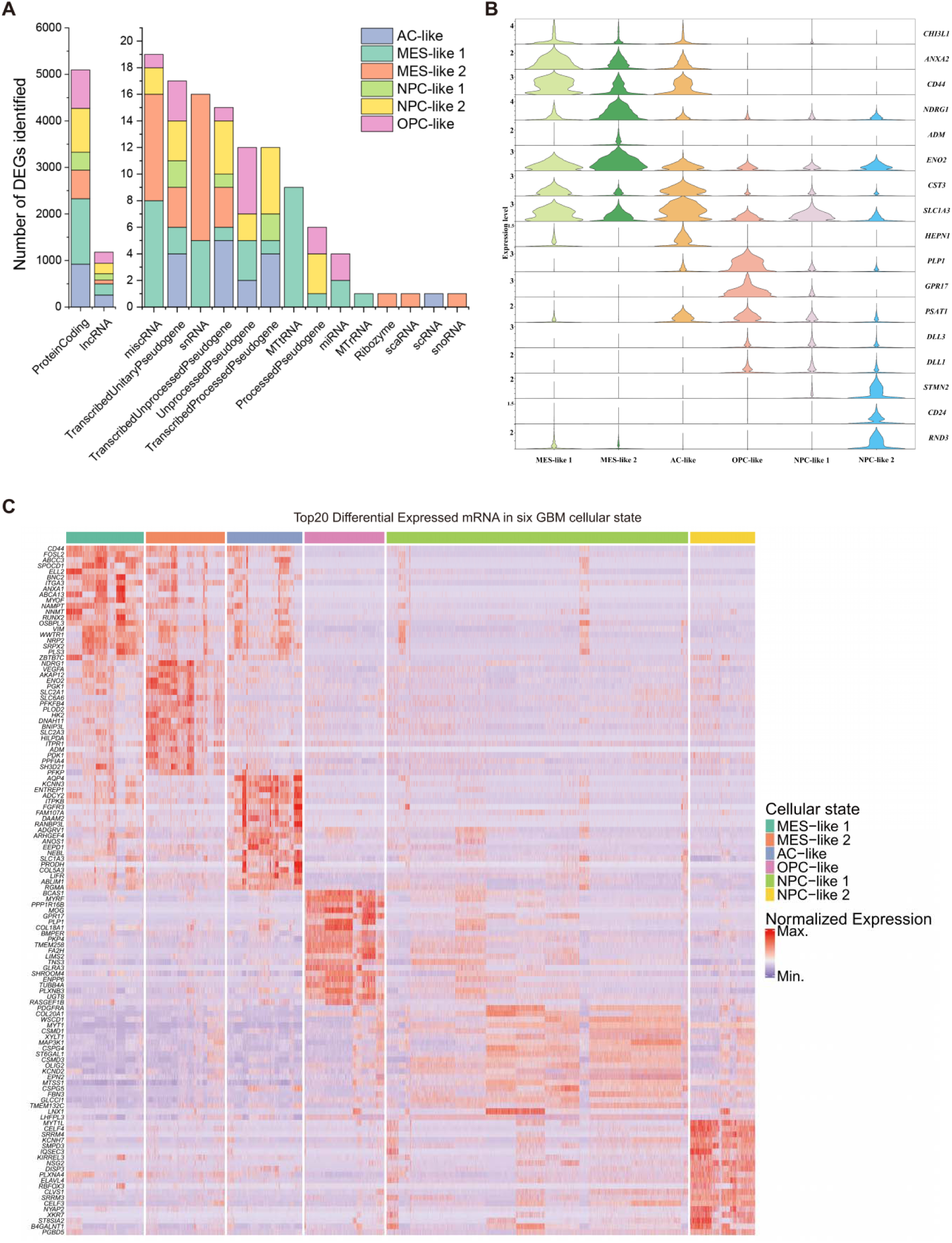
Differentially expressed genes across six glioblastoma cellular states. (A) Distribution of differentially expressed gene biotypes across six cellular states. (B) Violin plots of marker-gene expression across six cellular states. (C) Top 20 differentially expressed mRNAs across six cellular states.

**Extended Data Fig. 7.**
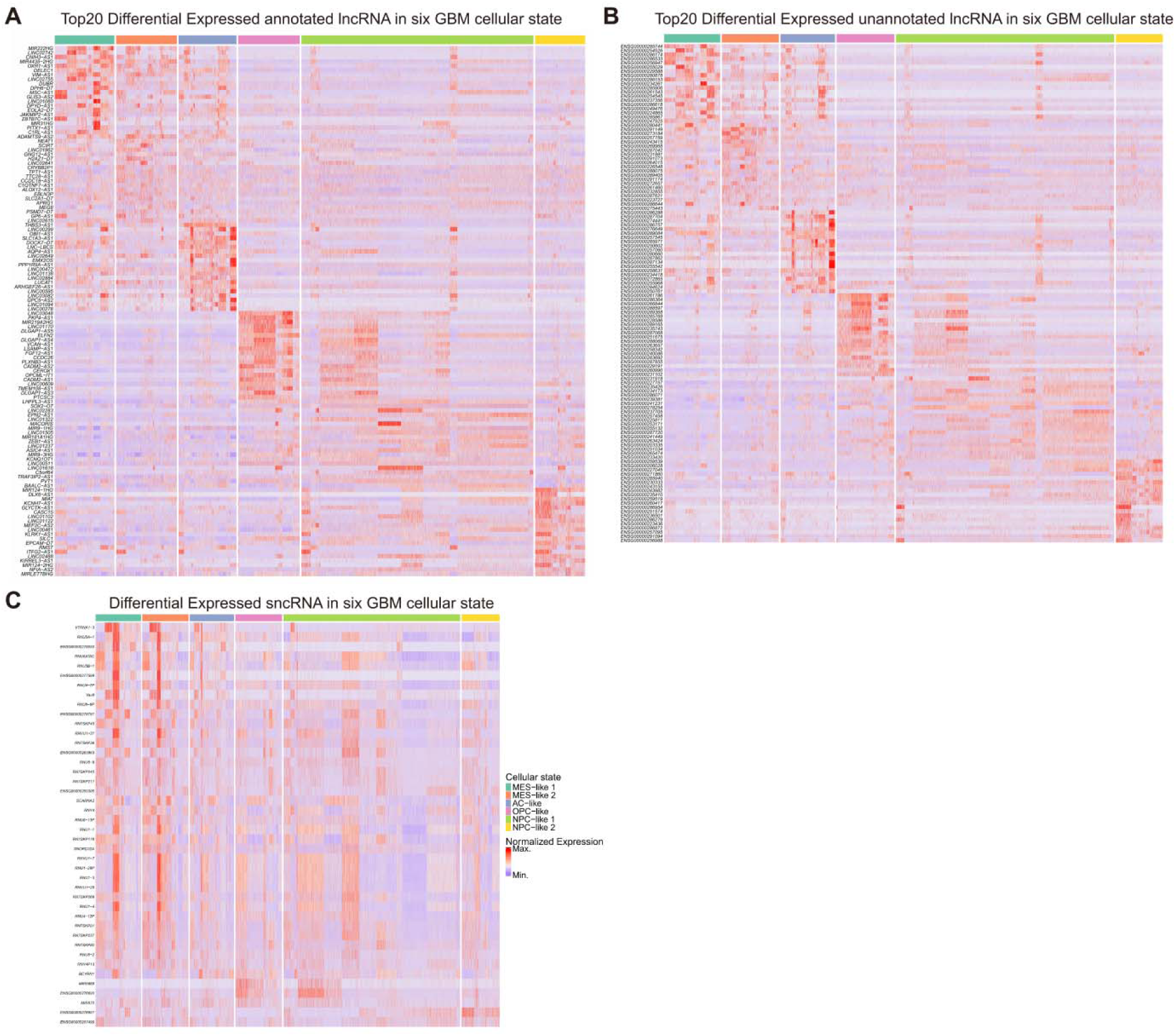
State-associated coding and non-coding RNA expression in glioblastoma.

**Extended Data Fig. 8.**
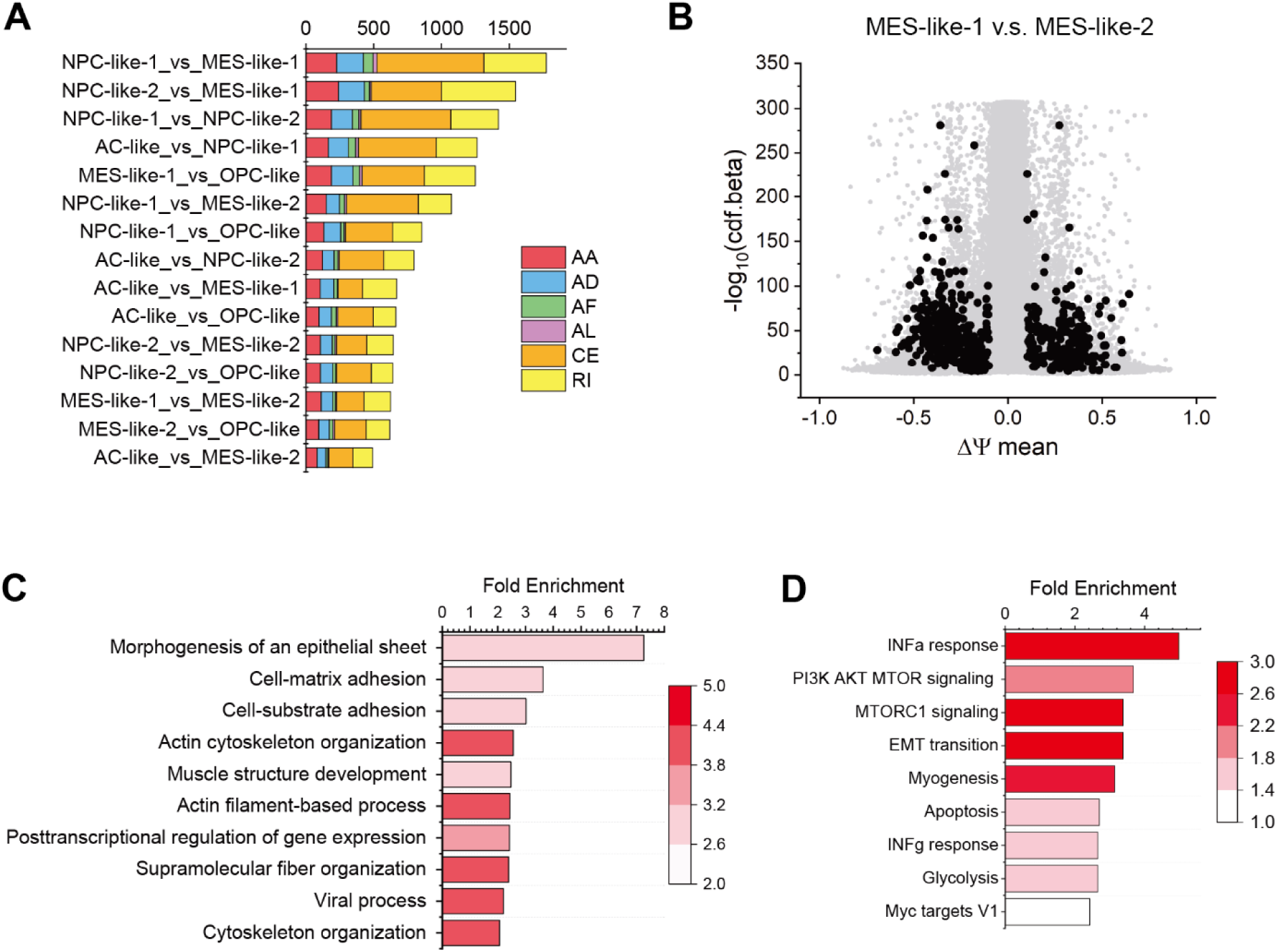
Differential splice-node inclusion across six glioblastoma cellular states. (A) Number of DISNs identified in all comparisons across 6 cellular states. b, Volcano plot of DISNs between the MES-like 1 and MES-like 2 states. (C and D) GO enrichments of genes with DISNs identified between MES-like-1 and MES-like-2 states using hallmark (C) and biological process GO terms (D).

**Extended Data Fig. 9.**
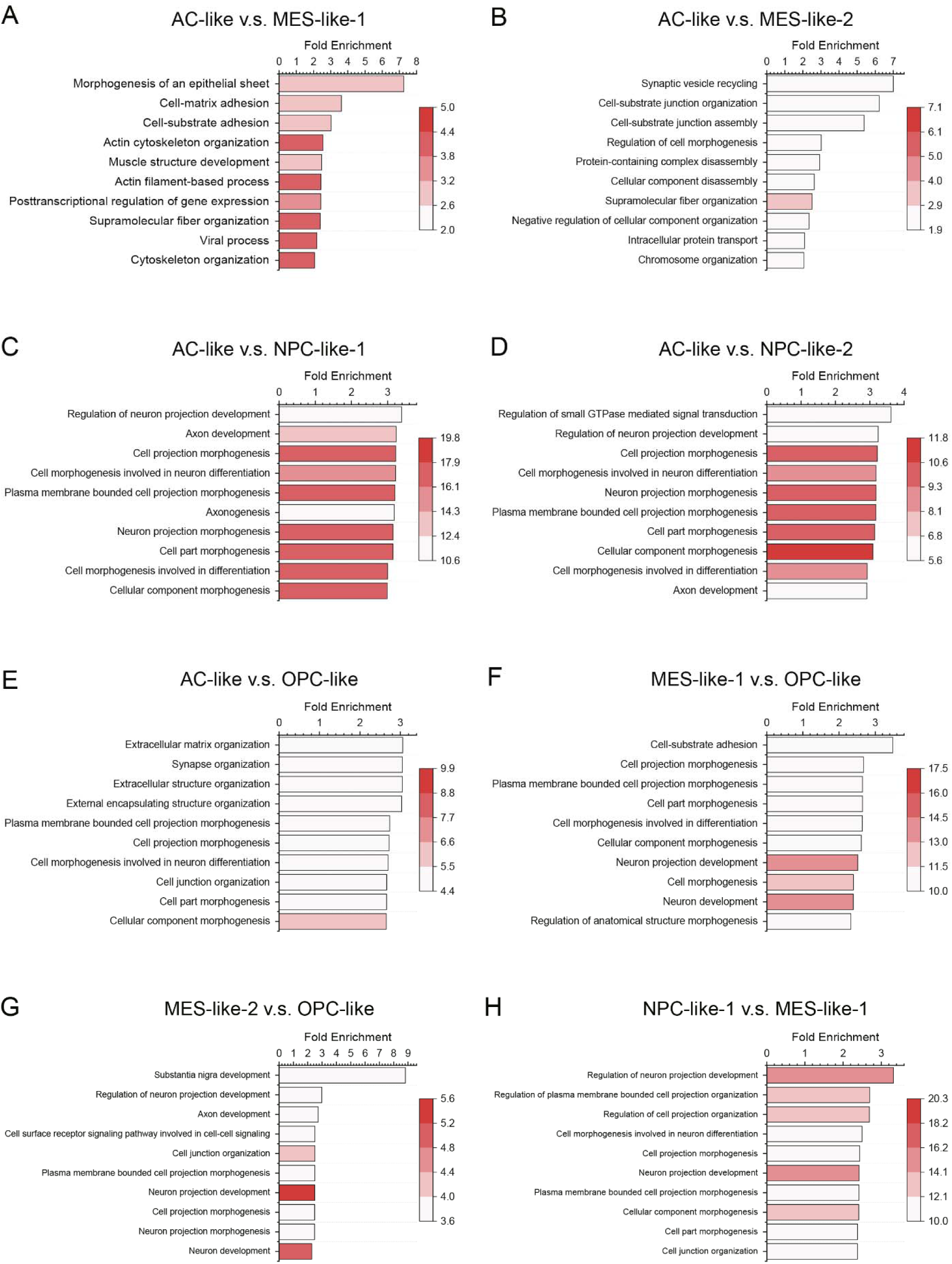
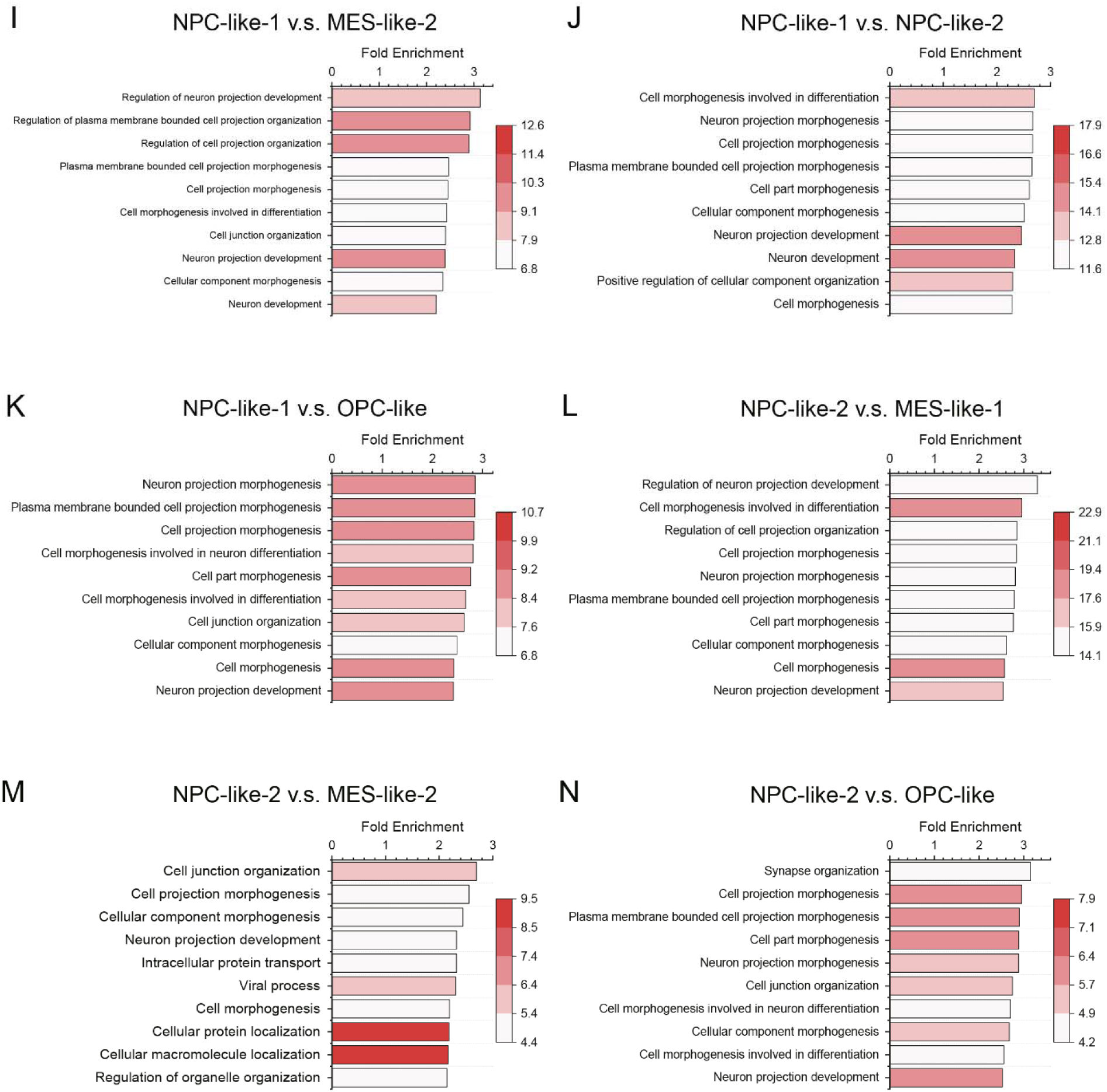
Functional enrichment of genes with differentially included splice nodes.

