## Supplemental Table 1 for "Decoding the Transcriptome Dark Matter: Construction of Single-Cell Whole-Transcriptome Regulatory Atlas by dropTotal"

^6^Lead contact

Table S1. ‌Patient Clinical Information

| Patient ID | State | Age/Sex | 2021 Classification | WHO Grade |
| --- | --- | --- | --- | --- |
| Patient 0 | Primary | 49/F | Astrocytome, IDH-mutant | 2 |
|  | Recurrent | 50/F | Glioblastoma, IDH-mutant | 4 |
| Patient 1 | Primary | 52/M | Glioblastoma, IDH-mutant | 4 |
|  | Recurrent | 53/M | NA | NA |
| Patient 2 | Primary | 52/M | Glioblastoma, IDH-mutant | 3-4 |
|  | Recurrent | 53/M | Glioblastoma, IDH-mutant | 4 |
| Patient 3 | Primary | 57/M | Glioblastoma, IDH -WT | 4 |
|  | Recurrent | 58/M | Glioblastoma, IDH -WT | 4 |
| Patient 4 | Primary | 43/M | Glioblastoma, IDH -WT | 4 |
|  | Recurrent | 44/M | Glioblastoma, IDH -WT | 4 |
| Patient 5 | Primary | 62/M | Glioblastoma, IDH -WT | 4 |
|  | Recurrent | 67/M | Glioblastoma, IDH -WT | 4 |
| Patient 6 | Primary | 49/F | Oligodendroglioma & 1 p/19 q codel | 2 |
|  | Recurrent | 56/F | Oligodendroglioma & 1 p/19 q codel | 3 |
| Patient 7 | Primary | 50/M | Glioma, IDH-mutant | 2 |
|  | Recurrent | 54/M | Glioma, IDH-mutant | 3 |
| Patient 8 | Primary | 38/M | Oligodendroglioma & 1 p/19 q codel | 3 |
|  | Recurrent | 41/M | Oligodendroglioma & 1 p/19 q codel | 3 |
| Patient 9 | Primary | 48/F | Glioblastoma, IDH -WT | 4 |
|  | Recurrent | 48/F | Glioblastoma, IDH -WT | 4 |
| Patient 10 | Primary | 49/M | Astrocytome | 3 |
|  | Recurrent | 49/M | Glioblastoma | 4 |
| Patient 11 | Primary | 66/F | Glioblastoma, IDH -WT | 4 |
|  | Recurrent | 67/F | Glioblastoma, IDH -WT | 4 |
| Patient 12 | Primary | 51/M | Glioblastoma, IDH -WT | 4 |
|  | Recurrent | 52/M | Glioblastoma, IDH -WT | 4 |
